# Quantification of metabolic activity from isotope tracing data using automated methodology

**DOI:** 10.1101/2024.02.25.581907

**Authors:** Shiyu Liu, Xiaojing Liu, Jason W. Locasale

## Abstract

Isotope tracing is a widely used technique to study metabolic activities by introducing heavy labeled nutrients into living cells and organisms. However, interpreting isotope tracing data is often heuristic, and application of automated methods using artificial intelligence is limited due to the paucity of evaluative knowledge. Our study developed a new pipeline that efficiently predicts metabolic activity in expansive metabolic networks and systematically quantifies flux uncertainty of traditional computational methods. We further developed an algorithm adept at significantly reducing this uncertainty, enabling robust evaluations of metabolic activity with limited data. Using this technology, we discovered highly reprogrammed mitochondria-cytosol exchange cycles in tumor tissue of patients, and observed similar metabolic patterns influenced by nutritional conditions in cancer cells. Thus, our refined methodology provides robust automated quantification of metabolism allowing for new insight into metabolic network activity.

## INTRODUCTION

Metabolism is the collection of interconnected chemical reactions that occur within living organisms, with its status determined by both molecule concentrations and the rates at which molecules are converted within this network. Isotope labeling, employing both stable and radioactive isotopes, is a prevalent method for directly analyzing reaction rates in physiologically-relevant settings^1^. Organisms or cells are fed or infused with isotopically labeled substrates, and mass spectrometry (MS) or nuclear magnetic resonance (NMR) is utilized to quantify isotope incorporation into metabolites of interest. A common technique among various stable-isotope methods is steady-state labeling. In this approach, isotope-labeled substrates are administered until the labeling ratio does not change, referred as steady-state, and information of reaction activities is encoded those steady-state labeling patterns^2^. The typical procedure is to manually interpret these data and draw conclusions about target reaction activities based on appropriate knowledge of metabolic pathway structure^3^. With advancements in MS technology, an increasing range of metabolites can be quantified across diverse biological systems^4^. However, this increase in measurable metabolites and their interconnections amplifies the complexity of labeling patterns, necessitating more complicated mathematical analyses to fully exploit the potential of advanced MS technologies in metabolic research.

There are several automated methodologies to decipher labeling patterns based on machine learning, such as metabolic flux analysis (MFA). Typical machine learning algorithms train predefined models with known data and generate predictions based on learned parameters. On the contrary, the MFA algorithm trains a preset metabolic network with labeling patterns derived from experimentally measured metabolites, and directly outputs trained parameters, that is, all fluxes in the network that best fit labeling patterns^3,5^. Despite its unique approach, current MFA methods often lack systematic evaluation and benchmarking, which is a standard practice in other machine learning algorithms^6^. One common assumption without sufficient evidence is that the training step necessarily finds the best set of fluxes to fit the isotope pattern, and that the best-fitted flux is the real flux in the biological system^7-9^. Notably, the significant variability observed in best-fit fluxes within these data, has been acknowledged in MFA results but remains largely unaddressed by existing methodologies^3,10^. These and other limitations constrain the utility of these automated methodologies and impede its wider application in metabolism research.

To systematically analyze and advance the automated methodology for metabolism analysis, we developed a novel package that can accurately resolve over 100 reaction rates from experimental isotope labeling data generated by MS, all within seconds of CPU time. By rigorously computing flux solutions aligned with experimental tracing data, we first elucidated the inherent variability present in results from the traditional methodology. We then proposed a protocol utilizing our novel package to accurately quantify this flux variability. Additionally, we developed an algorithm designed to substantially mitigate this variability, yielding accurate and reproducible results. Moreover, we assessed our methodology in the face of different limitations in data availability, quantifying the variability with various types of data. Finally, we conducted several new analyses of isotope tracing data and discovered new insights into cancer metabolism, thus showing that such technology greatly expands the capabilities of current isotope-tracing based approaches in metabolism research.

## RESULTS

### Assessment of current analysis methods in labeling experiments

In labeling experiments, uniformly labeled glucose (U-^13^C) is a preferred tracer due to its cost-effectiveness, stability, and ability to be traced in a vast array of metabolic pathways. Labeled glucose is either fed or infused to cultured cells or animals, and ^13^C will gradually replace ^12^C atoms in metabolites during metabolism to form the variety of isotopomers, molecules only differing in isotope atoms. These ^13^C isotopomers can be identified and quantified by MS or NMR, and their relative ratios, referred to as mass isotopomer distribution (MID), are then fed to automated analysis methods, such as MFA, to evaluate metabolic activities (Figure 1A).

**Figure 1.**
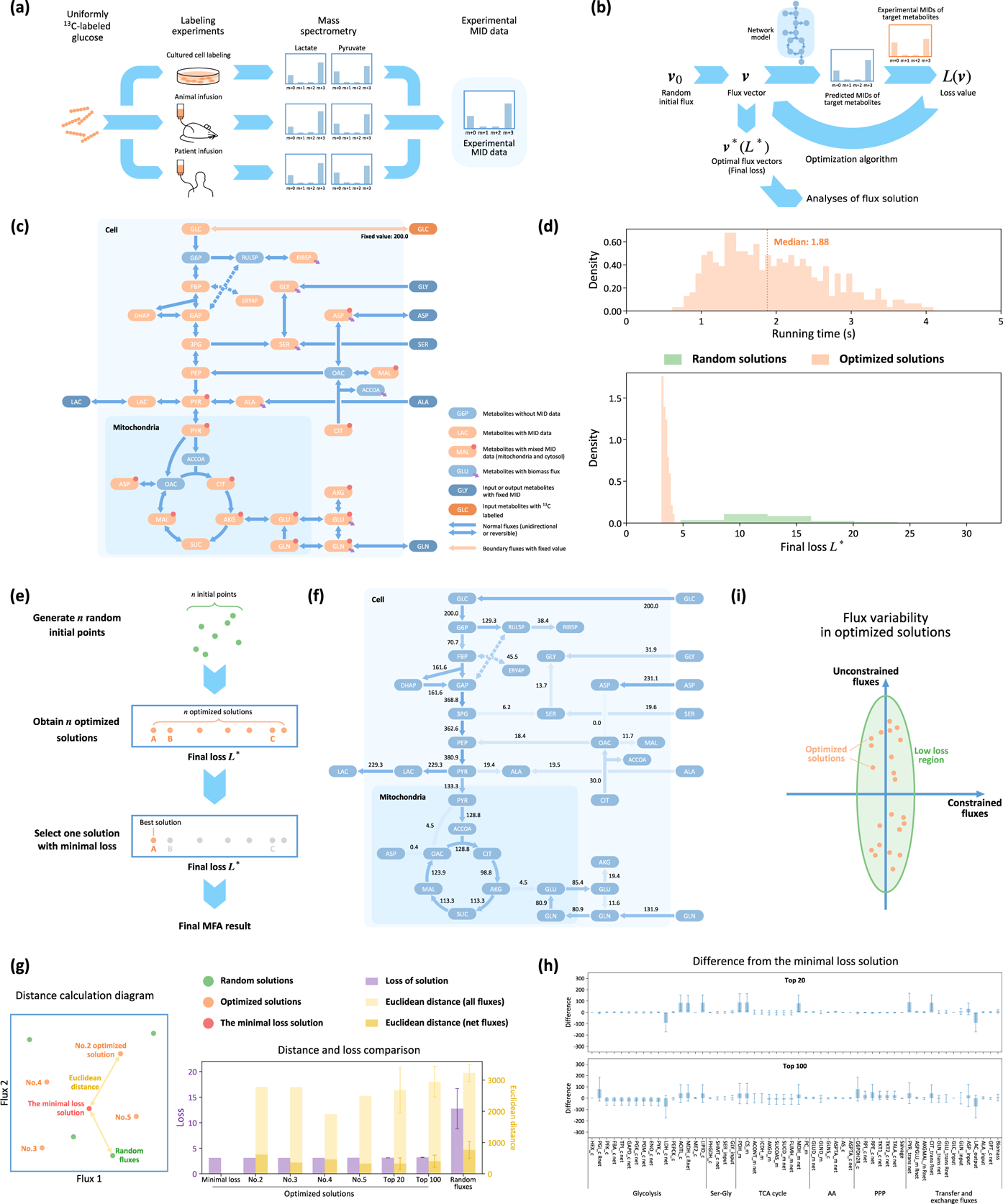
Comprehensive assessment of automated methodology using isotope-tracing data. (a) Data acquisition of mass isotopomer distribution (MID) data. (b) General steps of the prediction-optimization cycle in the metabolic flux analysis (MFA): The prediction step involves converting the flux vector to a loss function, while optimization step updates the flux vector based on the loss function and optimization algorithm. (c) Metabolic network model and data availability in the dataset of the cultured cancer cell line HCT-116. Note that not all metabolites and fluxes are plotted in this diagram. (d) Distribution of running time for optimized solutions (upper panel) and the final loss of 400 optimized and 10000 randomly obtained solutions (lower panel). The median running time is indicated by an orange dashed line in the upper panel. (e) Traditional method for generating and selecting the solution with the minimal loss from a set of optimized solutions. (f) Presentation of the flux analysis result from the HCT-116 dataset obtained using the traditional method, detailing the calculated flux values of the minimal-loss solution. (g) Comparison of loss values between top optimized solutions and randomly generated solutions, alongside their respective Euclidean distances from the minimal loss solution for all fluxes and net fluxes. Error bars represent standard deviations for each solution group. (h) Detailed examination of the net flux deviations in the top 20 and top 100 optimized solutions from the minimal loss solution, featuring standard deviation error bars for each respective group. (i) Illustration of flux variability in optimized solutions, indicating substantial variance in certain fluxes.

MFA utilizes MIDs to evaluate metabolic activities within a given metabolic network model, finding a flux vector that best aligns with measured MIDs as a solution. This process involves two primary steps: prediction and optimization. Starting from an initial guess of a flux vector, the prediction step calculates MIDs from the model and vector, determining the loss value, which is the difference between predicted and actual MID. The optimization step then updates the flux vector based on the loss value and the specific optimization algorithm, iteratively refining the vector until convergence (Figure 1B). In typical MFA applications, multiple solutions are generated from different initial points and the one with the minimal loss value is selected, thus avoiding local minima^8,9^.

To enhance the efficiency of MFA and broaden its application to complex metabolic networks and extensive isotope tracing data, we developed a new automated analysis package using Python. Its prediction step utilizes the commonly used elementary metabolite units (EMU) framework^11^, and the optimization algorithm is the sequential quadratic programming (SLSQP) in SciPy^12^. To verify ability of this package in large model, we constructed a metabolic network including several common pathways, such as glycolysis, the tricarboxylic acid (TCA) cycle, the pentose phosphate pathway (PPP), one-carbon metabolism, and several amino acid (AA) biosynthetic pathways. This model includes around 100 fluxes, much larger than what most flux measurements aim to compute. The model also features organelle compartmentalization, independently calculating the MID of one metabolite in the mitochondria and cytosol to approximate the real situation in eukaryotes (Figure 1C).

The model and package were tested with data from a uniformly glucose labeled (U-^13^C) cultured HCT116 cells^13^, which include MIDs covering most pathways in our network model (Figure 1C). Simulations starting from random initial points indicated that optimization processes typically finish within 2 seconds, yielding solutions with significantly lower and more consistent loss values compared to those obtained from initial points (Figure 1D, methods). The MIDs predicted from these optimized solutions also showed a closer match to experimental data than those predicted from the initial points (Figure S1A). The solution with minimal loss was then selected, which is the standard strategy utilized in current MFA methods (Figure 1E). The minimal loss solution found that around 60% of pyruvate is converted to lactate, 35% enters TCA cycle, and 5% is converted to alanine (Figure 1F). This metabolic pattern generally fits prior flux calculations on HCT116 cells^13^.

However, the conclusion becomes intricate when analyzing more solutions. We compared the optimized solutions with those randomly obtained solutions generated from a hit-and-run sampling algorithm^14^. Using principal component analysis (PCA) to project both optimized and randomly obtained solutions onto a two-dimensional plane, we observed that optimized solutions are more sparsely distributed and have greater mutual distances than the random solutions (Figure S1B). Interestingly, even the top-performing optimized solutions, although similar in loss value to the minimal loss solution, were found to be geometrically far from one another (Figure 1G). Further examination revealed that these top optimized solutions exhibit large variability in certain net fluxes, despite a close range of loss values (Figure 1H, S1C). This indicates that loss values alone are insufficient for estimating certain fluxes, leading to substantial uncertainties in optimized solutions (Figure 1I). Thus, the traditional method of selecting a single solution is inadequate for large-scale metabolic networks with a multitude of fluxes, underscoring the need for a new protocol that can consistently produce accurate solutions in such complex scenarios.

### Development and benchmark of a precise and robust algorithm for metabolic activity measurement

To assess accuracy of a solution, we considered simulated MID data that are predicted from a known flux vector (*υ*’). Accuracy can thus be evaluated by comparing the resulting flux vector with *υ*’ (Figure 2A). Given the observed high variability in certain fluxes among top optimized solutions (Figure 1H), we hypothesized that averaging solutions with minimal loss values would produce more accurate solutions. Therefore, we devised an optimization-averaging algorithm, that selecting *m* solutions with the minimal loss from *n* candidate optimized solutions (selected solutions) and averaging them to obtain new solutions (averaged solutions) (Figure 2B). When *m* equals to 1, the protocol reduces to the traditional method (Figure 1E). We also explored further optimization starting from these averaged solutions, producing what we termed re-optimized solutions (Figure 2B). The efficacy of selected, averaged and re-optimized solutions was assessed by their Euclidean distance from *υ*’ and the deviation in each net flux.

**Figure 2.**
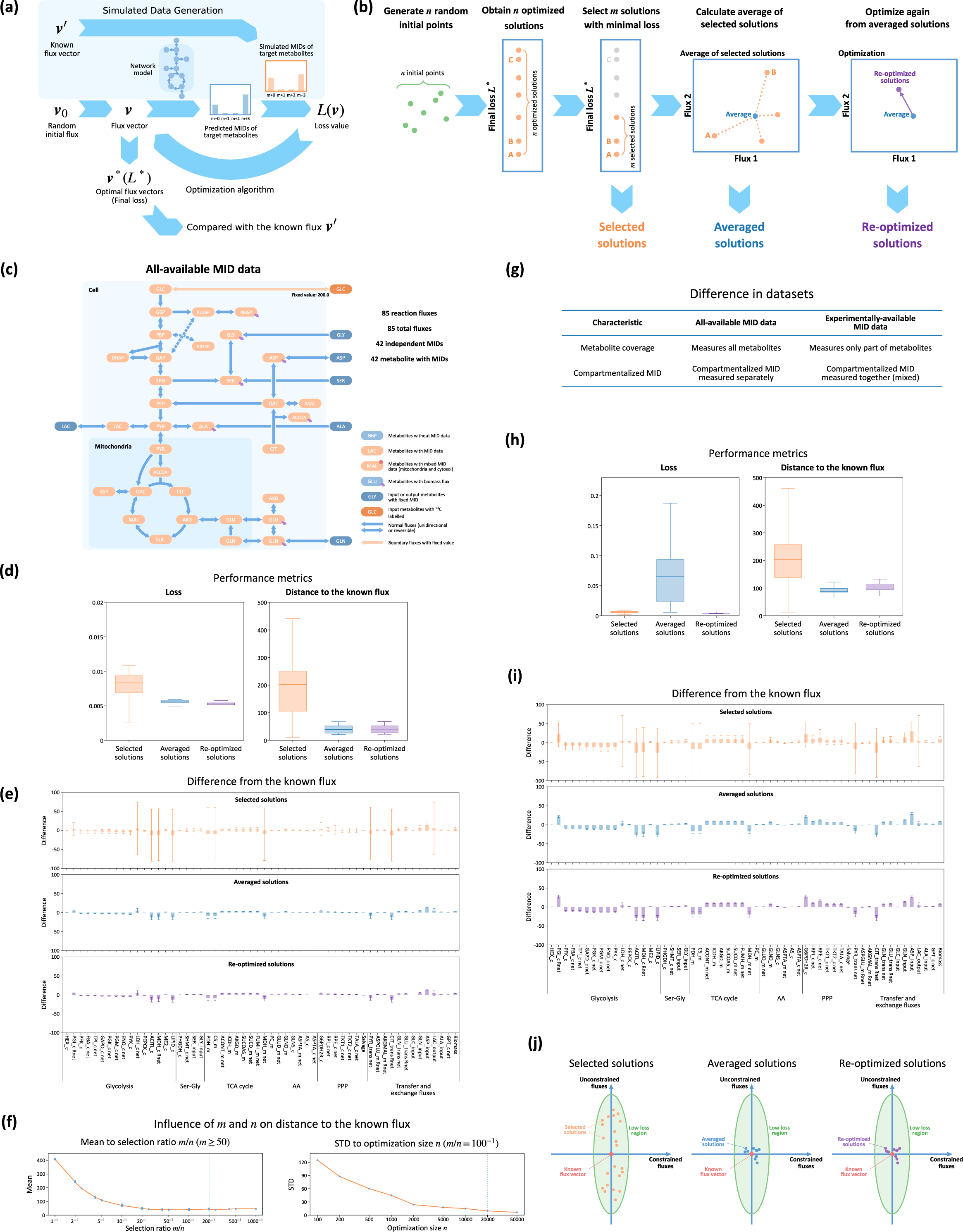
Development of the optimization-averaging algorithm. (a) General steps to generate simulated MID data from the known flux *υ*’ and to obtain optimized solution from this simulated data. (b) Diagram illustrating the optimization-averaging algorithm and the process of generating selected, averaged and re-optimized solutions. (c) Metabolic network model and data availability for the all-available dataset. (d, h) Comparison of loss values and the Euclidean distances of net fluxes from the known flux *υ*’ for selected, averaged and re-optimized solutions in the all-available dataset (d) and the experimentally-available dataset (h). Box plots represent quantile distributions, and whiskers indicate the full range of data. (e, i) Detailed examination of specific net flux deviations from the known flux *υ*’ in selected, averaged and re-optimized solutions within the all-available dataset (e) and the experimentally-available dataset (i), including standard deviation error bars for each group. (f) Changes in the mean Euclidean distances of net fluxes to *υ*’ relative to the selection ratio *m*/*n* (left panel) and alterations in the standard deviation (STD) of these distances in relation to the optimization number *n* (right panel) in averaged solutions. The x-axis of these line-plots is logarithmically scaled. (g) Illustration of the disparities between the all-available dataset and the experimentally-available dataset. (j) Diagram depicting the distribution of selected, averaged, and re-optimized solutions in the low loss region proximal to the known flux *υ*’.

Using our large-scale metabolic network model (Figure 1C), we assumed that all metabolites could be measured in compartmentalized organelles, creating an all-available MID dataset (Figure 2C). We found that averaged solutions exhibited similar loss values as selected solutions, but with substantially reduced mean and uncertainties of distances from the known flux (Figure 2D, S2A). Specific net fluxes exhibited high variability in selected solutions, including lactate dehydrogenase (LDH_c) and citrate synthase (CS_m), consistent with our prior findings in the HCT116 dataset (Figure 1H, S1C). These fluxes were estimated more accurately in averaged solutions, with relative errors under 10% (Figure 2E, S2B). Further optimization, leading to re-optimized solutions, did not notably alter these flux estimations nor significantly refine their accuracy (Figure 2D, 2E, S2A, S2B, S2C). The elevated precision and robustness in averaged solutions are validated across various distantly-distributed known fluxes (Figure S2D). Additionally, we found that reducing the *m*/*n* ratio improved the precision of averaged solutions, while increasing *n* enhanced their robustness, thereby validating our parameter choices in balancing precision and robustness within a feasible total running time (Figure 2F, S2E). In conclusion, this optimization-averaging algorithm facilitates the generation of robust and accurate solutions.

The preceding results were based on the assumption of full MID availability for all network metabolites. However, in practical applications, experimental data often only covers a subset of metabolites, with mixed compartmentalized MIDs. Therefore, the algorithm was evaluated with simulated MID data reflecting the limited availability typically in experimental settings, creating an experimentally-available dataset (Figure 2G, Table S1). In this scenario, the optimization-averaging algorithm consistently improved precision and robustness of solutions, effectively reducing deviations in specific net fluxes (Figure 2H, 2I). Despite further optimization reducing the loss values of averaged solutions, it did not consistently improve and sometimes decreased the precision in certain fluxes (Figure 2H, 2I, S2F, S2G, S2H). Like the all-available MID dataset, we evaluated the algorithm’s performance across a range of known flux vectors (Figure S2I) and explored the impact of different *m* and *n* parameters in this limited dataset (Figure S2J). These observations confirm that loss values alone are inadequate for accurate flux estimation in scenarios with limited MID data. However, our optimization-averaging algorithm effectively overcomes this challenge, producing consistently accurate results in line with those expected from typical isotope tracing experiments (Figure 2J).

### Robustness of the algorithm across varied data limitations

The availability of MID data, encompassing both quantity and distribution for each metabolite, varies in real isotope tracing experiments. To assess our algorithm’s performance under different data availabilities, we utilized simulated MID data to mimic scenarios with varying numbers of evenly-distributed available metabolites, pathway-specific MID exclusion, and compartmentalized MID measurements (Figure 3A, 3B, methods).

**Figure 3.**
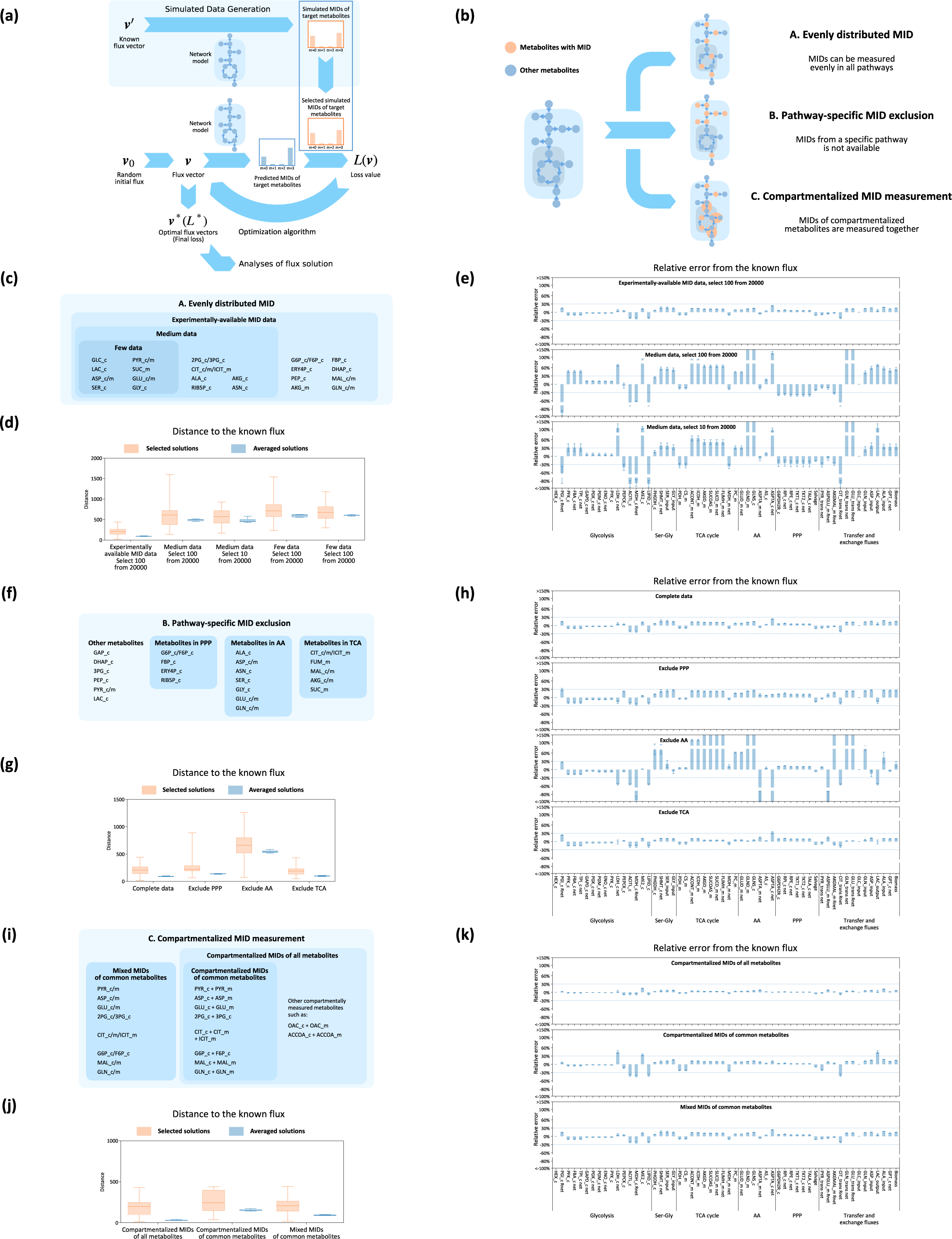
Robustness of the optimization-averaging algorithm under data limitations. (a) Outline of steps used to derive optimized solutions in the robustness analysis, with modifications in data availability highlighted in blue boxes. (b) Illustration of three typical scenarios in data availability: evenly distributed available metabolites, pathway-specific MID exclusion, and compartmentalized MID measurements. (c) List of measurable metabolites in three datasets with evenly distributed but varying number of MID data. (d, e) Comparison of the Euclidean distance of net fluxes (d) and the relative errors for each net flux (e) relative to the known flux in averaged solutions from evenly-distributed MID datasets. (f) List of measurable metabolites in three pathway-specific exclusion datasets designed to simulate the absence of all metabolites from specific metabolic pathways. (g, h) Comparison of the Euclidean distance of net fluxes (g) and relative errors for each net flux (h) relative to the known flux in averaged solutions from pathway-specific exclusion datasets. (i) Enumeration of measurable metabolites in three datasets assessing the impact of including or excluding compartmentalized MID measurements. (j, k) Comparison of the Euclidean distance of net fluxes (j) and relative errors for each net flux (k) relative to the known flux in averaged solutions from datasets with or without compartmentalized MID measurements. All box plots depict quantiles and whiskers show the range extremes. Error bars in all bar plots represent standard deviations.

In the scenario of evenly-distributed available metabolites, we examined cases with varying quantities of available metabolites, ranging from small to medium, and up to experimentally-available datasets (Figure 3C). Our optimization-averaging algorithm consistently reduced the mean distance and uncertainty to the known flux across all data sizes (Figure 3D), and effectively minimized deviation in each net flux (Figure 3E, S3A, S3B). However, as data availability decreased, the accuracy of estimated flux in averaged solutions also exhibited a significant decline (Figure 3E, S3G). Notably, selecting a smaller size *m* could partially compensate for increased fitting errors (Figure 3E). These results demonstrate that our algorithm, with more stringent selection criteria, can partially alleviate increased fitting errors associated with limited data availability.

We also explored the effects of excluding all metabolites from specific pathways, such as the PPP, AA biosynthesis, and TCA cycle, on the algorithm’s accuracy (Figure 3F). In these cases, our algorithm consistently demonstrated the ability to obtain high-precision estimated fluxes (Figure 3G, S3C, S3D). Notably, the precision of estimated fluxes exhibited particularly high sensitivity to the availability of AA biosynthetic pathways compared to the availability of PPP and TCA cycle (Figure 3H).

To determine the benefit of compartmentalized MID measurements, we created three different datasets (Figure 3I). Contrary to expectations, compartmentalized MIDs did not significantly enhance precision of results, showing that higher fitting precision in the all-available dataset was primarily due to the presence of more MID data (Figure 3J, 3K, S3E, S3F). This suggests that non-compartmentalized MIDs are generally adequate for precise flux analysis.

### Advanced automated methodology delineates the importance of exchange fluxes between mitochondria and cytosol in cancer patients

Exchange fluxes between mitochondria and cytosol, such as citrate/malate antiporter (CMA) and α-ketoglutarate/malate antiporter (AMA), are crucial in energy production and closely linked to specific metabolic patterns in cancer metabolism and cell state transitions^15,16^. However, these fluxes have been understudied due to difficulties in measuring them physiologically with traditional methods. Our advanced methodology is suitable for this condition and are utilized to analyze several published labeling data from patients with cancer that were infused with ^13^C-glucose^17,18^.

We selected a published dataset containing ^13^C-glucose-labeled kidney tumor and adjacent normal kidney tissue to analyze how metabolism in carcinoma is different from that in adjacent normal tissue *in situ* (Figure 4A). Our algorithm, using the same optimization-averaging algorithm and parameters as before (Figure 1B, 4B, 4C), and with a medium-available dataset (Figure 3C, 4D), converged to provide flux sets for each patient (Figure S4A, S4B).

**Figure 4.**
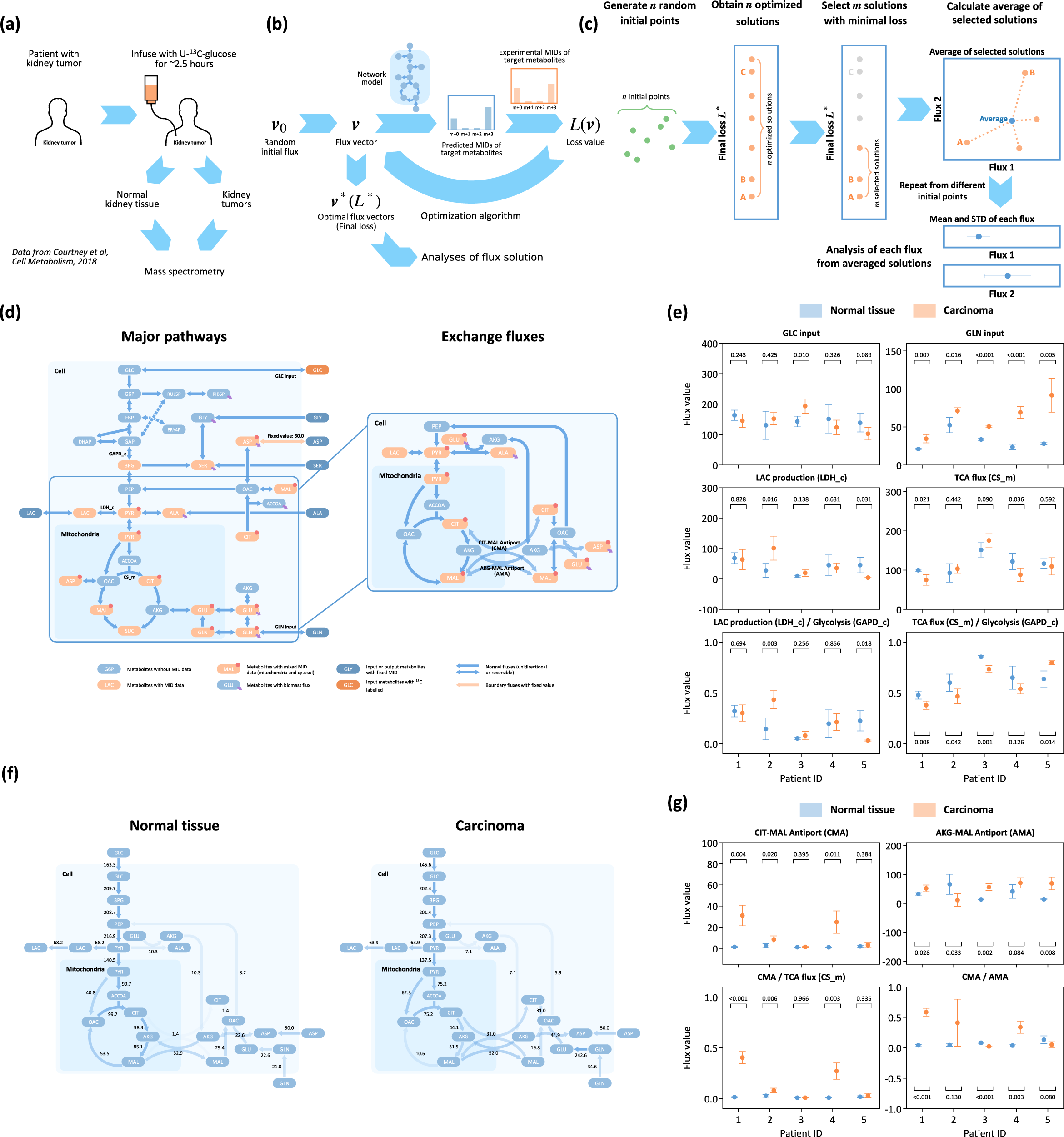
Analysis of *in vivo* labeling data from normal tissue and cancer in patients through the optimization-averaging algorithm. (a) Procedure of data acquisition: Normal tissue and cancer are obtained from the same patient who received an infusion of U-^13^C-glucose for 2.5 hours. The labeling patterns in these samples are then quantified by MS. (b) Outline of steps followed to derive optimized solutions from this experimental data. (c) Application of the optimization-averaging algorithm in this experimental context, detailing the calculation of final fluxes and their associated uncertainties. (d) Metabolic network model and data availability for this analysis. The right panel provides a zoomed-in view of a section of the metabolic network, highlighting the fluxes in TCA cycle and some exchange fluxes between mitochondria and cytosol. (e, g) Comparison of glycolytic and TCA fluxes (e) and mitochondrial-cytosolic exchange fluxes (g) between normal tissue and cancer. Data points (circles) and error bars represent the mean and standard deviation of the averaged solutions, respectively. Statistical significance (p-values) comparing normal tissue and carcinoma for each patient is presented in the figures. (f) Network map illustrating the major exchange fluxes between normal tissue and cancer in a representative patient (Patient 1). The map highlights the activation of the citrate-malate antiporter (CMA) in carcinoma. The numbers denote the mean flux values from the averaged solutions.

The results indicated that while glucose uptake and TCA cycle flux were similar in carcinoma and normal tissue, the ratio of TCA cycle flux to glycolytic flux decreased in three of the five patients (p-value ≤ 0.05, Figure 4E, S4C). A notable increase in glutamine input flux was also observed in all patients (p-value ≤ 0.05, Figure 4E, S4C). These variations confirmed that the Warburg effect, characterized by a larger fraction of glucose flux being directed to lactate rather than the TCA cycle, exists *in vivo* with precise flux results^17^. Compared to traditional analyses that rely on averaged isotope tracing data, our methodology allows for individual patient analysis to generate specific glycolytic and TCA fluxes, uncovering greater heterogeneity and enhancing clarity and interpretation of results.

A previous study has reported that CMA between mitochondria and cytosol are elevated during stem cell differentiation^16^. It is hypothesized that differential activation of CMA may reflect a change of cellular metabolic state. However, traditional methods are unable to accurately quantify these exchange fluxes involving multiple metabolites across different organelle compartments. Our tool provides an opportunity to study exchange fluxes in the background of the large-scale metabolic network under physiological condition. In a network diagram of patient 1, TCA cycle fluxes tend to leave mitochondria after citrate (CIT) by CMA in carcinoma, rather than to be converted to mitochondrial α-ketoglutarate (AKG) as normal TCA cycle (Figure 4F), which causes increased CMA flux and its ratio to TCA cycle fluxes (Figure 4G). This pattern was similarly noted in patient 4, underscoring its potential significance (Figure 4G). Conversely, the AMA flux remains relatively stable in carcinoma, resulting in a significantly increased CMA to AMA flux ratio in patient 1 and 4 (p-value ≤ 0.01, Figure 4G), highlighting a selective alteration in exchange fluxes.

Reanalysis of the data with the traditional approach of selecting a single solution with minimal loss from optimized solutions (Figure 1E) revealed significant flux uncertainty, obscuring the metabolic distinctions between tumor and normal tissues (Figure S4D, S4E). This highlights the superiority of our methodology in detecting nuanced variations in metabolic fluxes, particularly in peripheral pathways.

In addition to metabolic variations between cancer and normal tissue, tissue origin may also affect cancer metabolism. Our approach facilitates the analysis of metabolic differences across various cancer types simultaneously. By analyzing published datasets with kidney, lung and brain tumors from different patients with same model and protocol (Figure 4C, 4D, S5A), we observed distinct patterns: kidney tumors exhibited a stronger Warburg effect, with a higher lactate production to glycolytic flux ratio and a lower TCA cycle flux to glycolytic flux ratio compared to lung and brain tumors (Figure S5B). This observed differential Warburg effect is consistent with previous study, suggesting tissue origin plays a crucial role in cancer metabolism^17^. Additionally, the ratio of CMA to TCA cycle flux was notably higher in kidney tumors, underscoring the potential of this metric to reflect distinct metabolic states in different cancers (Figure S5C).

### Enhanced automated methodology unveils nutrient-dependent differential exchange fluxes in cultured cells

Nutritional microenvironment and availability have significant influences on cancer generation and development^19,20^. Although *in vivo* data from carcinoma patients can reflect real physiological state, analyzing the metabolic response of cancer cells to nutritional environment remains challenging. To further test our automated methodology and to investigate Warburg effect and exchange fluxes under varied nutritional conditions, we conducted tracing experiments and metabolic analysis with eight colon cancer cell lines (SW620, SW480, HCT8, HT29, HCT116, NCI, SW48, SW948). These cell lines were incubated in RPMI media supplemented with ^13^C-labeled high glucose concentration (11.1 mM) and low glucose concentration (0.5 mM) for 24 hours. MID data collected by MS (Figure 5A) were analyzed using our established metabolic network and optimization-averaging algorithm (Figure 5B, Figure 4C). Our tool effectively fitted almost all experimental MIDs, except for a few metabolites poorly detected by MS (Figure S6A, S6B).

**Figure 5.**
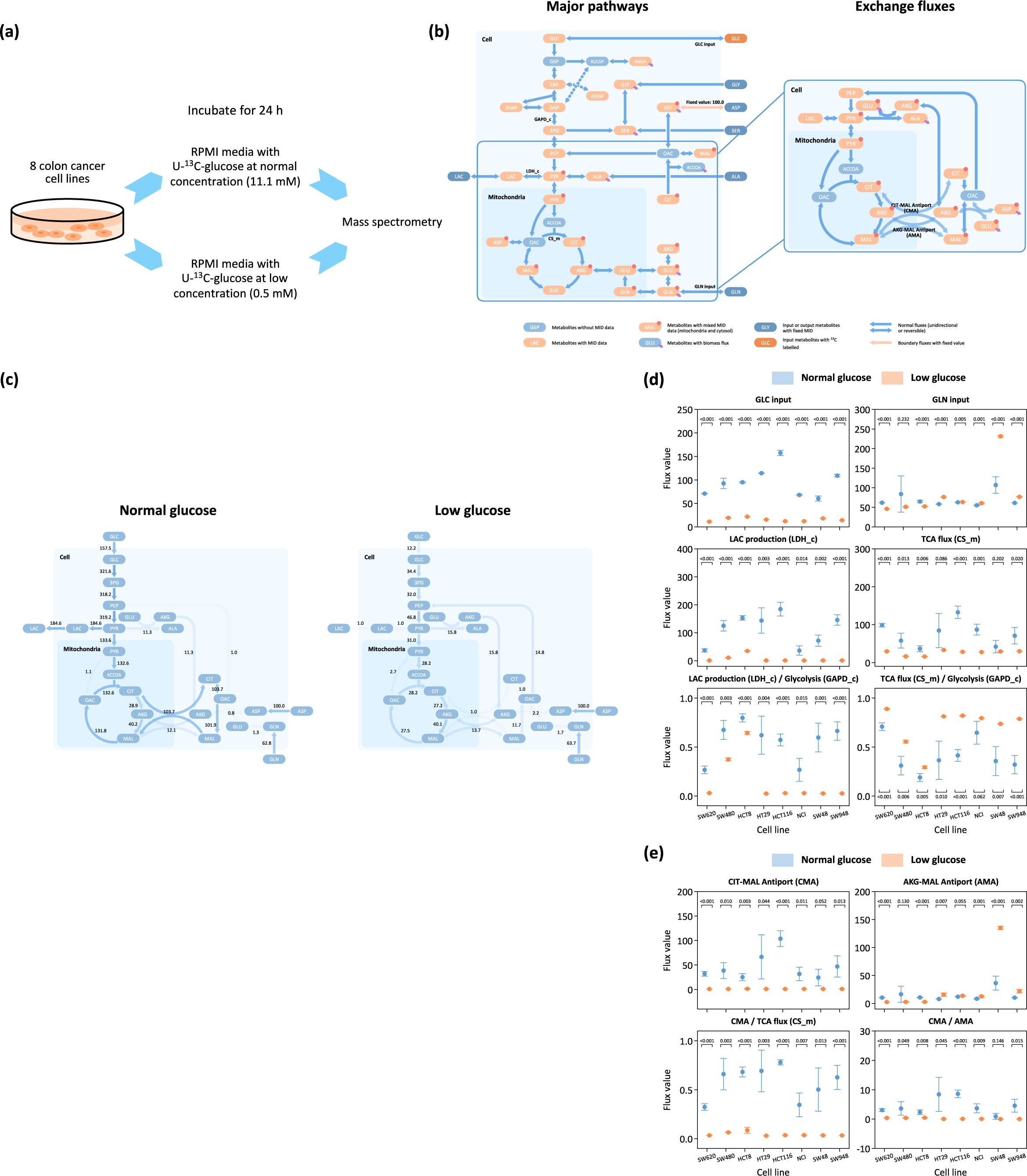
Analysis of labeling data from cultured cell lines through the optimization-averaging algorithm. (a) Procedure of data acquisition: Eight different colon cancer cell lines are incubated with normal and limited U-^13^C-glucose for 24 hours, and their labeling ratio is measured using MS. (b) Metabolic network model and data availability for this analysis. The right panel provides a zoomed-in view of a section of metabolic network, highlighting the fluxes in the TCA cycle and some exchange fluxes between mitochondria and cytosol. (c) Network map illustrating the major exchange fluxes under normal and limited nutrition in a representative cell line (HCT-116). The map highlights the active state citrate-malate antiporter (CMA) under normal conditions and inactive state in limited nutrition. The numbers denote the mean flux values from the averaged solutions. (d-e) Comparison of glycolytic and TCA fluxes (d) and exchange fluxes (e) under normal and limited nutrition in eight cancer cell lines. Data points (circles) and error bars represent the mean and standard deviation of the averaged solutions, respectively. Statistical significance (p-values) between normal and limited nutrition condition for each cell line is presented in the figures.

Our results indicate significant reductions in glucose input, TCA flux, and lactate flux under low-glucose conditions across all cell lines, reflecting a nutrient-deprived state (Figure 5C, 5D, S6C). Notably, lactate flux decreased more rapidly than glucose input, leading to a reduced lactate production to glycolytic flux ratio and an increased TCA cycle flux to glycolytic flux ratio under low-glucose conditions (Figure 5C, 5D, S6C). This pattern aligns with previous findings that link the Warburg effect to cellular nutrition and proliferative states^15^. As for exchange fluxes, CMA activity was prominent in high-glucose conditions but nearly absent in low-glucose environments (Figure 5E). This decrease in CMA flux, combined with the relative increase or stability of AMA flux, led to reduced CMA to TCA cycle flux ratio and CMA to AMA flux ratio (Figure 5C, 5E). Furthermore, we observed that the exchange flux patterns in cultured cancer cell lines resembled those in carcinoma patients, with nutrient limitation patterns resembling those in normal tissues (Figure 4F, 5C). This finding, aligned with prior studies on CMA variations during stem cell differentiation, suggests exchange fluxes might serve as indicators of overall nutritional and proliferative states of a cell^16^.

Contrastingly, analyses using the traditional method exhibited significant flux variability, undermining the clarity of metabolic differences under different nutrition conditions (Figure S6D, S6E). This highlights the precision and robustness of our advanced methodology, which excels in elucidating these complex metabolic variations.

## DISCUSSION

Our study developed a new methodology that is first built on a comprehensive assessment of the issues encountered during automated methods for isotope tracing data. Traditionally, the evaluation of metabolic activities has heavily depended on intuitive interpretations. Our automated method, enhanced by innovative algorithms, accurately calculates fluxes from not only complete datasets but also limited data. These advances address key concerns about the precision and robustness of automated methodologies, thereby broadening their applicability in diverse metabolic studies. Furthermore, our assessments and benchmarks offer fresh perspectives on analysis of metabolic activities, highlighting issues such as the high variability of fluxes in solutions with closely clustered loss values (Figure 1G) and the discrepancy between loss values and the actual distance of net fluxes from the known flux in certain solutions (Figure 2H). Additionally, our findings question the assumed benefit of compartmentalized data in enhancing the precision of net flux estimates in models with multiple compartments (Figure 3K). These insights encourage a reevaluation of the information extracted from MID data and suggest a need to refine experimental approaches to optimize the trade-off between cost and utility in isotope tracing studies.

In representative applications, our methodology has proven effective in quantifying key metabolic fluxes, notably mitochondria-cytosol exchange fluxes, which have recently been recognized for their importance in key cellular processes. These fluxes play a pivotal role in the malate-aspartate shuttle (MAS), crucial for maintaining NAD+ levels and the redox balance between the mitochondria and cytosol^21^. By accurately quantifying these fluxes under physiological conditions, our approach enhances understanding of redox and energy homeostasis within biological systems, offering insights into, for example, aging and autoimmune diseases^22,23^. Moreover, integrating this methodology with additional omics, physiological, and spatial data promises to unveil new discoveries. For instance, pinpointing biomarkers indicative of high metabolic flux states could significantly advance our comprehension of health, aging, and disease, areas where current biomarkers are notably insufficient.

Despite the advancements our algorithm offers in mitigating precision loss due to data limitations, the scarcity of available MID measurements remains a significant challenge. This scarcity directly impacts the accuracy, precision, and overall effectiveness of flux analyses. The quality of isotope tracing experiments, indicated by the range of measurable metabolites, the accuracy of MS quantifications, and the variability in batched data, remains a cornerstone for precise flux estimation and metabolic research. The complexity of conducting tracing experiments in living organisms often exacerbates these challenges, limiting the broader application of *in vivo* flux analysis. Enhancements in experimental methodologies and data acquisition techniques are crucial to fully exploiting the capabilities of our advanced automated flux analysis approach.

## METHODS

### Data Sources

This study is based on two data sources: the *in vivo* labeling data were obtained from infused patients in previous work^17,18^, while the labeling data of cultured cell lines were acquired using methods based on previous studies ^24,25^.

### Metabolic flux analysis

### Flux model and constraints

The flux model includes around 100 reactions including central carbon metabolism, major branch pathways, and two compartments (Figure 1C). All metabolites, except input and output metabolites labeled in figure, were assumed to be balanced during simulations, which means sum of income fluxes to a certain metabolite equals to sum of outgo fluxes from this metabolite.

### MID prediction and flux optimization

Given a set of flux values, MID of target metabolites can be predicted from averaging MID of precursors using flux values as weights:

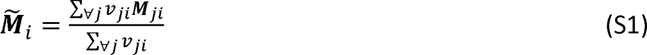

where 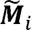 is the predicted MID vector of metabolite *i*, ***M**_ji_* is MID of metabolite *i* produced from a substrate *j*, and *υ_ji_* is the flux from *j* to *i*.

Some metabolites exist both in mitochondria and cytosol, and therefore extra independent pool size variable of each compartment for those metabolites is introduced, and their mixed MID is calculated by average weighted by their pool size.

The difference between the predicted (possibly mixed) and experimental MIDs was evaluated by Kullback–Leibler divergence:

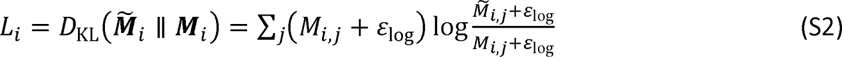

which *M_i,j_* and 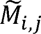 are element *j* in vector ***M**_i_* and 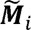 respectively. *ε*_log_ is a small number added to maintain numerical stability. Sum of *L_i_* for all target metabolites was regarded as the loss function *L*_total_ to minimize. The optimization problem was defined as:

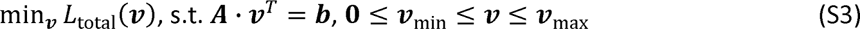

where ***A* · *υ***^*T*^ = ***b*** represents flux balance requirement and other equality constraints. ***υ***_min_ and ***υ***_max_ are lower and upper bounds for composite vector *υ*. This optimization problem is solved by SLSQP algorithm in SciPy package^12^.

### Random solution generation

Random solutions are used as initial feasible solutions in SLSQP algorithm, and as a negative control to study the distribution of optimized solutions. Feasible solutions are defined by all solutions that fit the same constraints that ***A*** · ***υ****^T^* = ***b*** and **0** ≤ ***υ***_min_ ≤ ***υ*** ≤ ***υ***_max_ as Eq. S3. A previously developed hit-and-run(HR)-derived algorithm named optGpSampler^14^ was adapted to this study to sample randomly distributed feasible solutions.

### Traditional MFA strategy

The traditional MFA method, as depicted in Figure 1E, involves running *n* independent optimizations from randomly obtained initial points. The solution with the lowest loss is selected as the final result. For consistency in this study, *n* is set at 400 across all traditional MFA applications, aligning with typical flux analysis scales.

For the analyses of real datasets shown in Figure S4D, S4E, S6D and S6E, we replicated this procedure 5 times, each with different a set of *n* randomly obtained points. The 5 selected solutions with the minimal loss from each set were combined to form the final flux analysis result for that dataset.

### Flux embedding

In section “Labeling experiments and automated analysis method”, random and optimized solutions are combined and embedded to two-dimensional space by principal component analysis (PCA) with the function sklearn.decomposition.PCA in Python package scikit-learn.

### Development of optimization-averaging algorithm

The development of our optimization-averaging algorithm, depicted in Figure 2B, involves conducting independent *n* optimizations. Each optimization begins with a randomly generated initial point, progressing to an optimized result using SLSQP algorithm. Among those optimized results, *m* solutions with the smallest loss values are selected (selected solutions) and are then averaged to form a single solution (averaged solutions). The averaged solutions may undergo further optimization to yield re-optimized solutions.

This optimization-averaging algorithm is repeated 30 times starting from different initial solution sets. For each repetition, distance (all fluxes and net fluxes) from the selected, averaged and re-optimized solutions to the known flux are calculated. Additionally, detailed differences and relative errors for all net fluxes in these solutions are also calculated with parameters *n* = 20000 and *m* = 100 except specifically labeled. Final figures include results from all 30 repetitions (Figure 2D, 2E, 2H, 2I, S2A, S2B, S2E, S2F).

To validate our algorithm with distant known flux vectors, we generated 30 known flux vectors, each having a net distance greater than 500 from one another (Figure S2C, S2G). For each of these vectors, the optimization-averaging algorithm was applied 3 times with parameters *n* = 20000 and *m* 100. The resulting selected, averaged, and re-optimized solutions were then subjected to similar analyses for validation.

To assess the impact of parameters *m* and *n* on the algorithm’s performance, we repeated the entire optimization-averaging process 30 times for each combination of *m* and *n*. The resulting solutions— selected, averaged, and re-optimized—were collected for analogous analyses (Figures 2F, S2D, S2H). To illustrate the precision trend, we plotted the mean and standard deviation (STD) of the distance (net fluxes) to different series of *m* and *n* values (Figure 2F).

### Robustness to data limitation analysis

In robustness to data limitation analysis, raw simulated data are first generated based on the known flux *υ*’. This data is then subjected to variations in data availability, simulating limited data scenarios. These adjusted datasets undergo our standard optimization-averaging algorithm, typically set with parameters *n* = 20000 and *m* = 100 except specifically labeled. The resulting selected and averaged solutions were collected, and their net distances to the known flux, as well as detailed differences and relative errors for all net fluxes, are analyzed as previously.

### Experimental data analysis

In this study, the MIDs from all biological replicates were averaged to minimize batch preparation and measurement errors, creating a target MID for the automated methodology. The optimization-averaging algorithm (*n* = 20000 and *m* = 100) was then applied to these experimental datasets, generating averaged solutions. This algorithm was repeated 5 times, and the resulting 5 averaged solutions were used as the final results for flux analysis of this dataset.

### Visualization of metabolic network map

In all figures, network diagrams with or without flux value are plotted with Python package Matplotlib. For clarity, not all reactions are displayed in one network diagram. In network diagram with flux value, directions of a flux arrow are set by directions of net flux and transparency of flux arrows are set based on its value.

### Software implementation

Scripts in this study were implemented with Python 3.8. Source codes are available from GitHub (https://github.com/LocasaleLab/Automated-MFA-2023). The package version dependency is also provided on GitHub website.

### Manuscript preparation

In preparing our manuscript, we utilized OpenAI’s ChatGPT 4.0 for text editing. This involved a series of iterative revisions with ChatGPT’s suggestions to refine language to improve readability. All revisions, including the final manuscript, underwent careful review by our team to ensure its accuracy.

## ACKNOWLEDGEMENTS

We express our sincere appreciation to the members of the Locasale laboratory for their valuable discussions. Special thanks are extended to Yudong Sun, Jared Rutter, and Arion Kennedy for help with editing the manuscript. Support from National Institutes of Health (R01CA193256) and the American Cancer Society (RSG-16-214-01-TBE) are gratefully acknowledged.

## AUTHOR CONTRIBUTIONS

S.L., and J.W.L. designed the study and wrote the manuscript. S.L. designed the algorithm, developed the package and performed the data analysis. X. L. performed labeling experiments of cultured cells and collected all experimental data. All authors have read, edited and approved the final manuscript.

---

**Figure S1.**
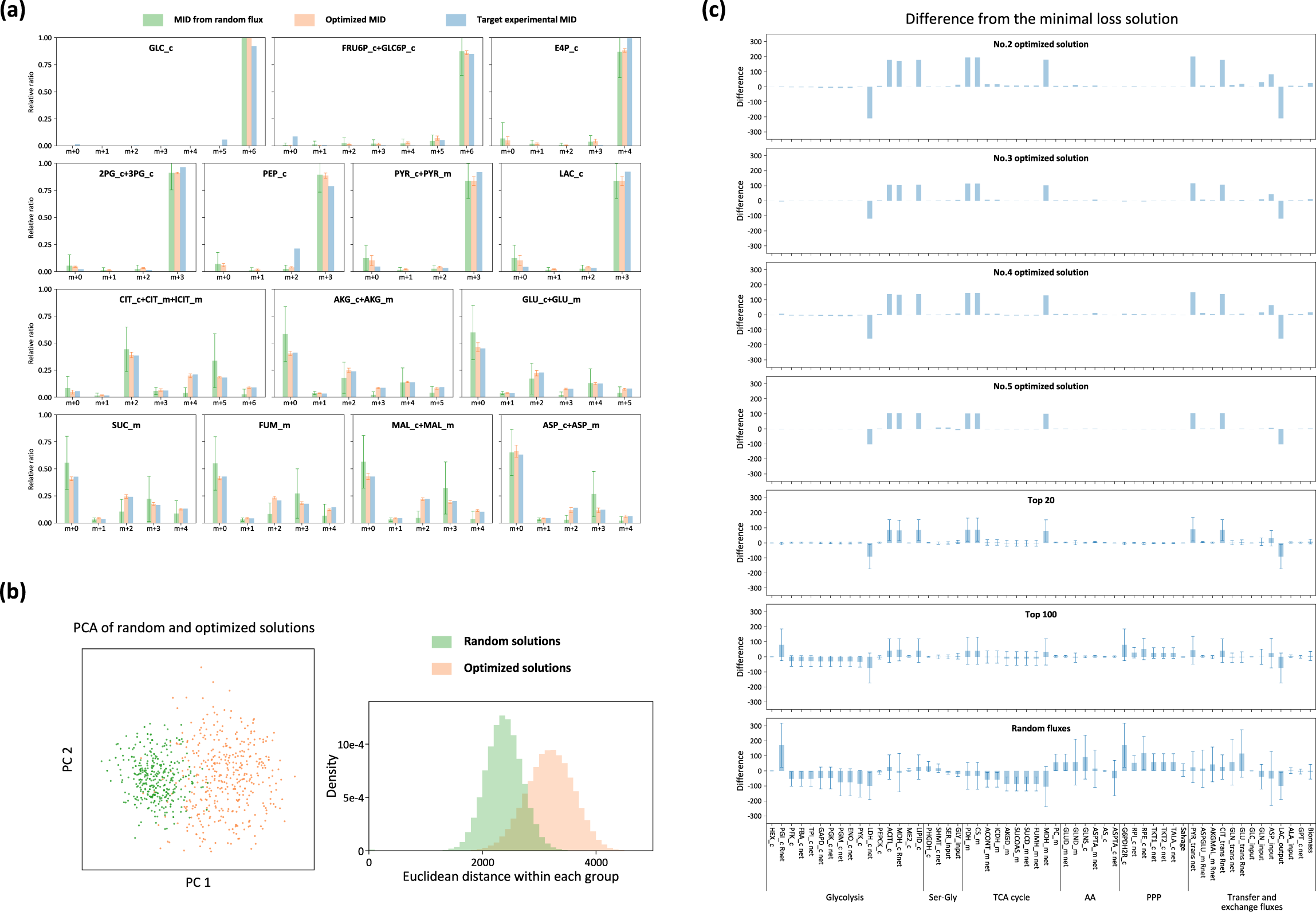
Details of the assessment of automated methodology. (a) Comparison of MID of several metabolites between randomly obtained solutions, optimized solutions and the experimental data. Error bars represent the standard deviation within both the randomly obtained and optimized solution sets. (b) Visualization of optimized and randomly obtained solutions using PCA (left) and distribution of mutual distance of optimized solutions and randomly obtained solutions (right). (c) Detailed examination of the net flux differences in various top-tier optimized solutions (including individual, top 20, and top 100 solutions) in comparison to the optimal solution, featuring standard deviation error bars for each respective group.

**Figure S2.**
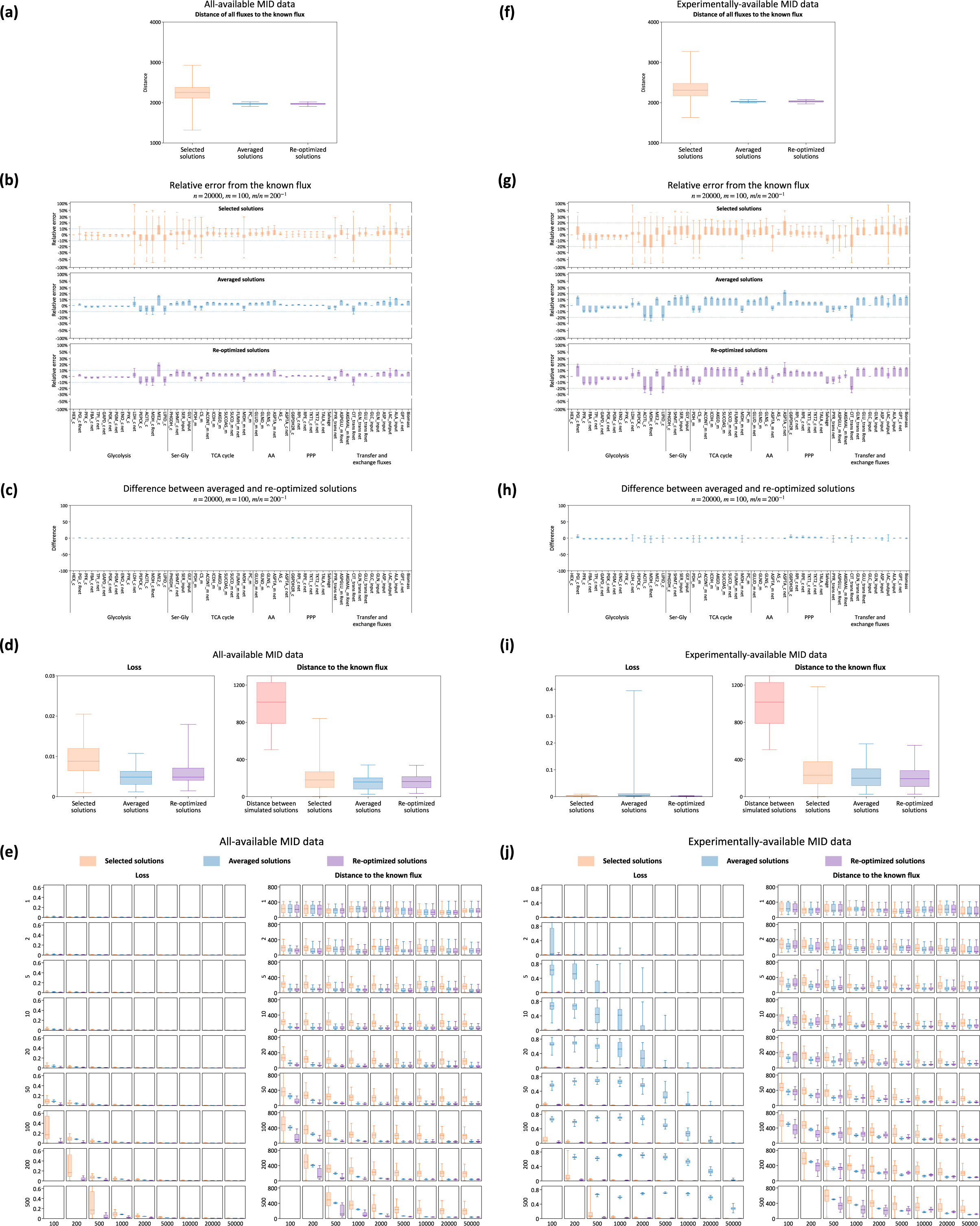
Details of the development of the optimization-averaging algorithm. (a, f) Comparison of distance of all fluxes from the known flux for selected, averaged and re-optimized solutions. The analysis includes both the all-available dataset (a) and the experimentally-available dataset (f). (b, g) Detailed examination of the net flux relative error compared to the known flux in selected, averaged and re-optimized solutions. The analysis includes both the all-available dataset (b) and the experimentally-available dataset (g). Each group includes standard deviation error bars. A blue dashed line represents a ±10% error range in (b) and ±20% in (g), serving as a reference for comparison of flux accuracy. (c, h) Detailed examination of specific net flux differences between each averaged solution and corresponding re-optimized solution, including standard deviation error bars, within the all-available dataset (c) and the experimentally-available dataset (h). (d, i) Comparison of loss values, mutual distance among 30 distant known fluxes, and distance of net fluxes from selected, averaged and re-optimized solutions to these known fluxes. The analysis includes both the all-available dataset (d) and the experimentally-available dataset (i). All optimized solutions displayed substantially lower distances to the known fluxes, compared to the mutual distances among known fluxes themselves. (e, j) Impact of different combinations of parameters *m* and *n* on the loss and distance of net fluxes from the known flux for selected, averaged and re-optimized solutions. This analysis includes both the all-available dataset (e) and the experimentally-available dataset (j). X-axis represents parameter *n* and y-axis represents parameter *m*.

**Figure S3.**
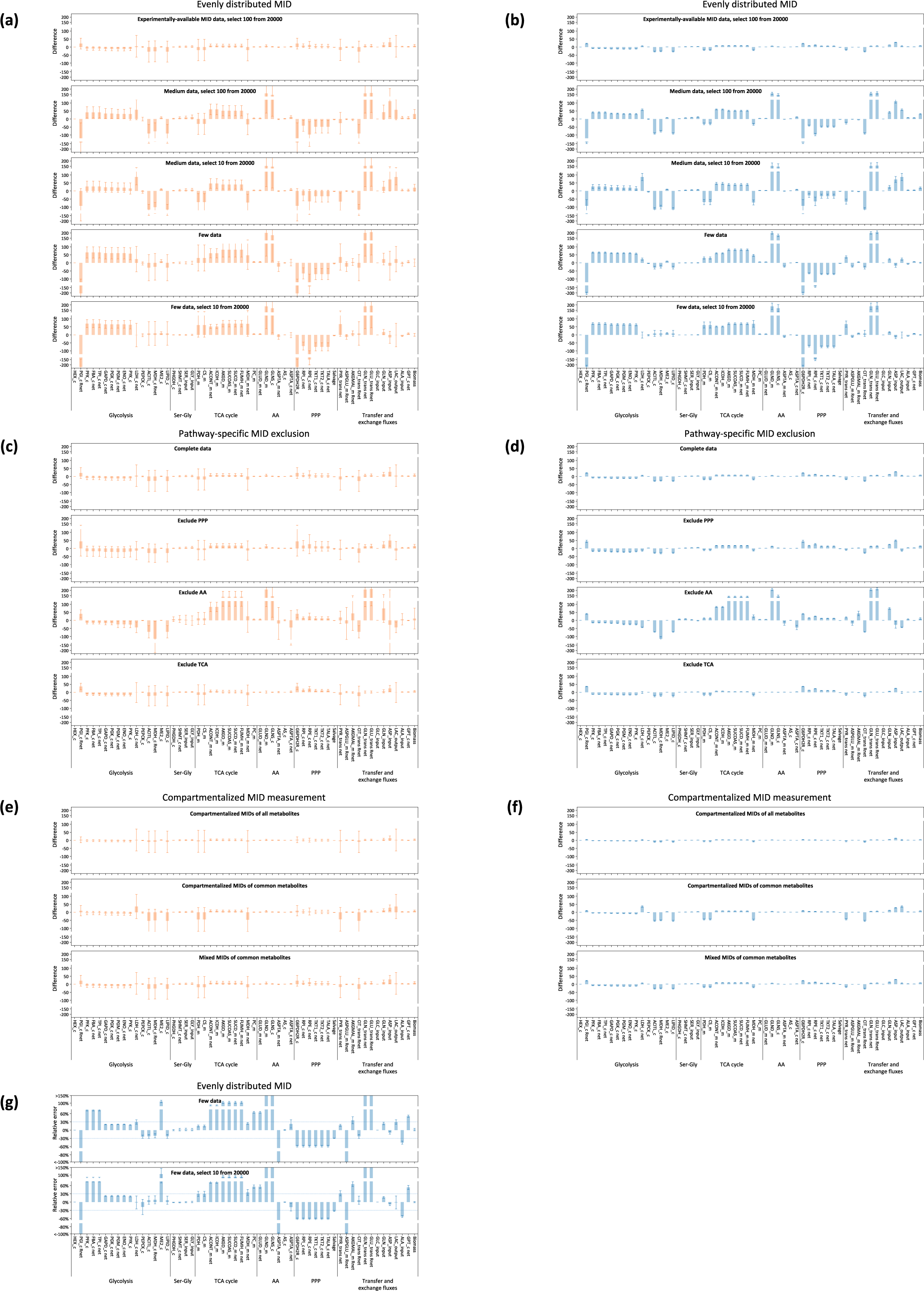
Details of robustness analysis of the optimization-averaging algorithm under data limitations. (a-f) Detailed examination of the deviations of each net flux relative to the known flux across different data limitation contexts, including evenly-distributed MID datasets (a, b), pathway-specific exclusion datasets (c, d), and datasets with or without compartmentalized MID measurements (e, f). This analysis covers both selected solutions (a, c, e) and averaged (b, d, f) solutions. (g) Comparison of the relative error for each net flux compared to the known flux in averaged solutions across two evenly distributed MID datasets. Error bars in all bar plots represent standard deviations.

**Figure S4.**
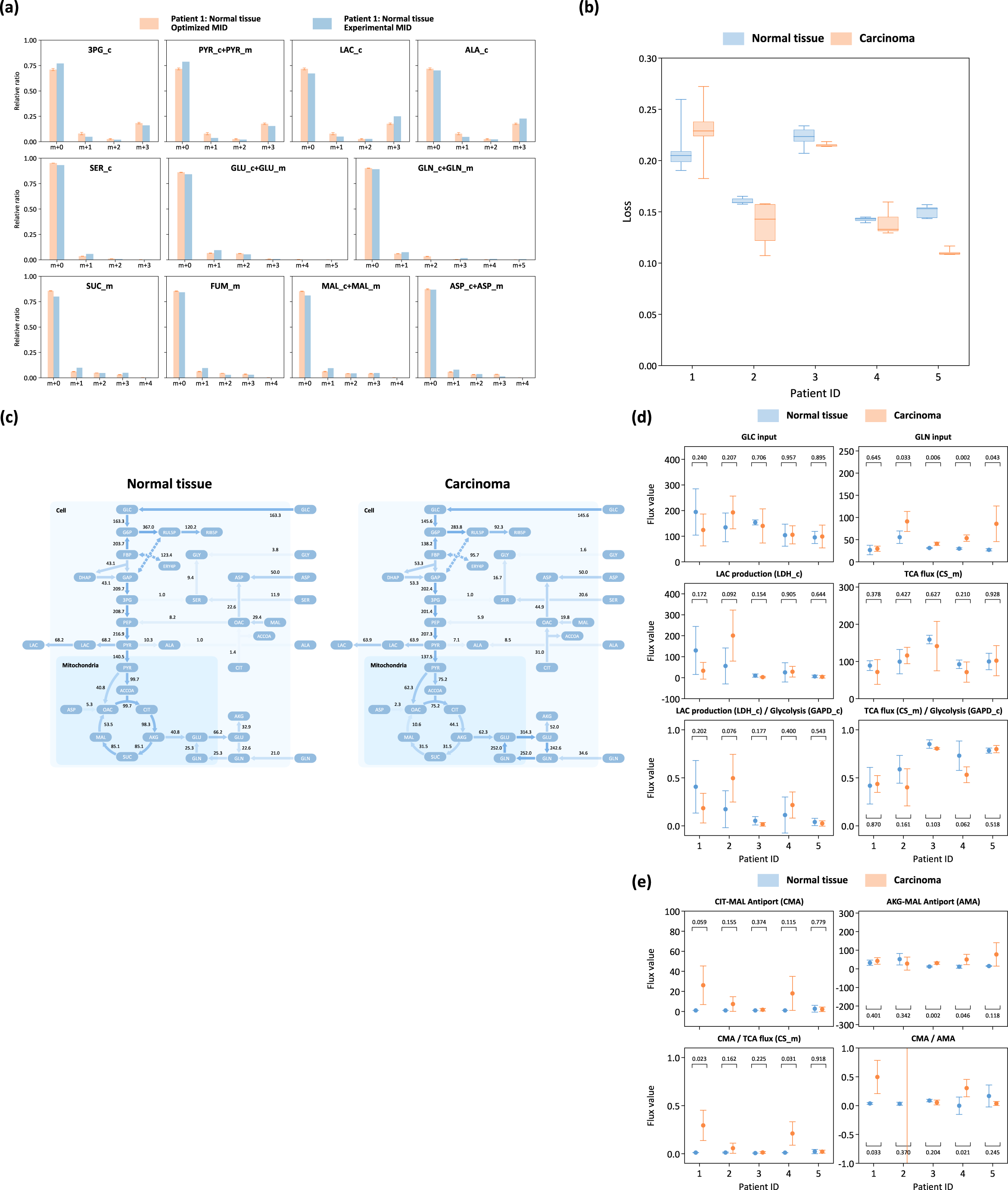
Details for the analysis of *in vivo* labeling data in patients with cancer through the optimization-averaging algorithm. (a) Comparison between the MIDs of all averaged solutions and experimental data in the normal tissue of the patient 1. Error bars represent the standard deviation of MID. (b) The distribution of the final loss values in all averaged solutions from normal tissue and cancer datasets are depicted across all five patients. (c) The complete network map of the major pathways of normal tissue and cancer in patient 1. (d-e) Comparison of glycolytic and TCA fluxes (d) and mitochondrial-cytosolic exchange fluxes (e) between normal tissue and cancer, calculating with the traditional method. Data points (circles) and error bars represent the mean and standard deviation of the averaged solutions, respectively. Statistical significance (p-values) comparing normal tissue and carcinoma for each patient is presented in the figures.

**Figure S5.**
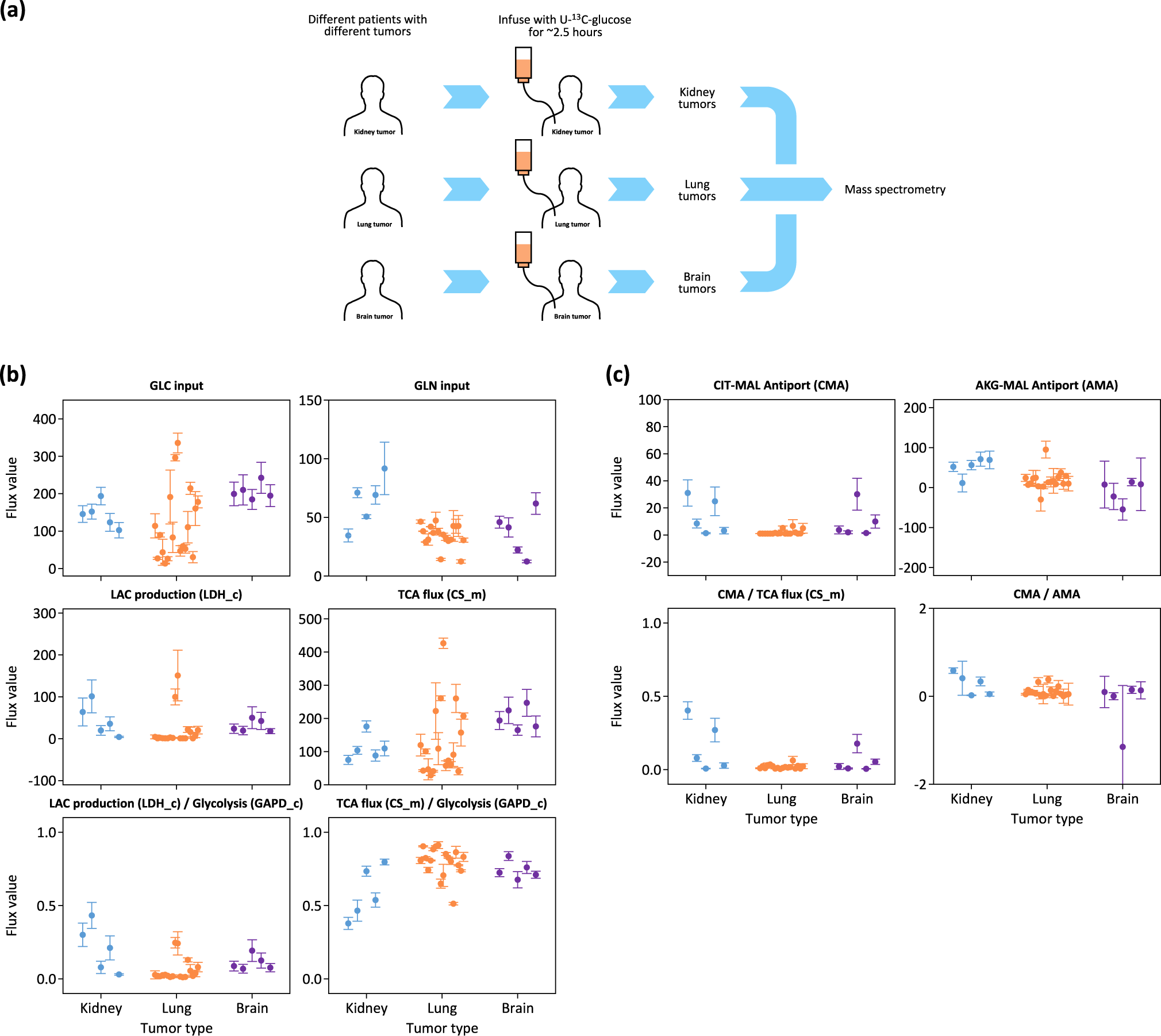
Analysis of *in vivo* labeling data obtained from multiple types of cancer across different patients through the optimization-averaging algorithm. (a) Description of the data acquisition procedure involving sampling and measurement of kidney tumor, lung tumor and brain tumor from patients infused by U-^13^C-glucose for 2.5 hours, using MS. (b-c) Comparison of glycolytic and TCA fluxes (b), as well as exchange fluxes (c), among kidney, lung and brain tumors. The sample from each patient is analyzed and displayed independently. Data points (circles) and error bars represent the mean and standard deviation of the averaged solutions, respectively.

**Figure S6.**
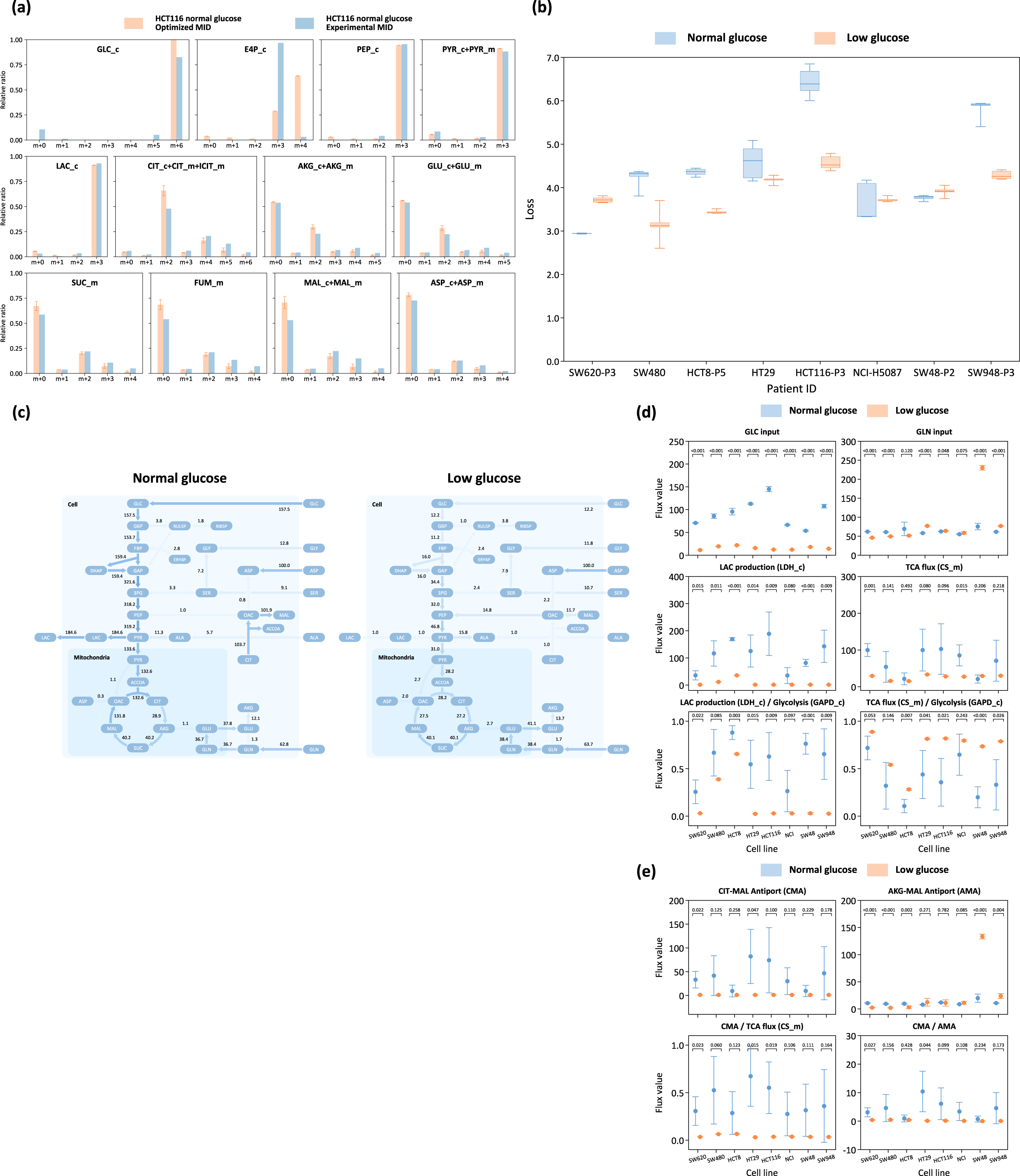
Details for the analysis of labeling data from cultured cell lines through the optimization-averaging algorithm. (a) Comparison between the MIDs of all averaged solutions and experimental data for HCT-116 cell line under normal conditions. Error bars represent the standard deviation of MID. (b) The distribution of the final loss values of in all averaged solutions from datasets under normal and limited nutrition conditions are depicted across all eight cell lines. (c) The complete network map of the major pathways of HCT-116 cell under normal and limited nutrition conditions. (d-e) Comparison of glycolytic and TCA fluxes (d) and mitochondrial-cytosolic exchange fluxes (e) under normal and limited nutrition conditions in eight cancer cell lines, calculating with the traditional method. Data points (circles) and error bars represent the mean and standard deviation of the averaged solutions, respectively. Statistical significance (p-values) between normal and limited nutrition conditions for each cell line is presented in the figures.

**Table S1.** Comparison of MID availability in all-available and experimentally-available MID datasets. For each metabolite, it is measured in cytosol (c), mitochondria (m), or both compartments combined (c+m). Metabolites measured as mixtures, like citrate and isocitrate, are also noted.

| All-available MID data |  | Experimentally-available MID data |  |
| --- | --- | --- | --- |
| Metabolite name | Compartment | Metabolite name | Compartment |
| glutamate | c | glutamate | c+m |
| glutamate | m |  |  |
| erythrose 4-phosphate | c | erythrose 4-phosphate | c |
| lactate | c | lactate | c |
| glycine | c | glycine | c |
| dihydroxyacetone phosphate | c | dihydroxyacetone phosphate | c |
| phosphoenolpyruvate | c | phosphoenolpyruvate | c |
| glucose 6-phosphate | c | fructose 6-phosphate/glucose 6-phosphate | c |
| fructose 6-phosphate | c |  |  |
| succinate | m | succinate | m |
| pyruvate | c | pyruvate | c+m |
| pyruvate | m |  |  |
| malate | c | malate | c+m |
| malate | m |  |  |
| glucose | c | glucose | c |
| 2-phosphoglycerate | c | 2-phosphoglycerate/3-phosphoglycerate | c |
| 3-phosphoglycerate | c |  |  |
| fumarate | m | fumarate | m |
| asparagine | c | asparagine | c |
| ribulose 5-phosphate | c | ribose 5-phosphate | c |
| glutamine | c | glutamine | c+m |
| glutamine | m |  |  |
| citrate | c | citrate/isocitrate | citrate(c+m)+isocitrate(m) |
| citrate | m |  |  |
| isocitrate | m |  |  |
| alanine | c | alanine | c |
| aspartate | c | aspartate | c+m |
| aspartate | m |  |  |
| a-ketoglutarate | c | a-ketoglutarate | c+m |
| a-ketoglutarate | m |  |  |
| serine | c | serine | c |
| fructose 1,6-bisphosphate | c | fructose 1,6-bisphosphate | c |
| oxaloacetate | c |  |  |
| oxaloacetate | m |  |  |
| ribose 5-phosphate | c |  |  |
| sedoheptulose 7-phosphate | c |  |  |
| acetyl-coa | c |  |  |
| succinyl-coa | m |  |  |
| acetyl-coa | m |  |  |
| 13BPG | c |  |  |
| glyceraldehyde 3-phosphate | c |  |  |
| xylose 5-phosphate | c |  |  |

