## Supplementary Methods for "Quantification of metabolic activity from isotope tracing data using automated methodology"

### General Method

#### Flux model

In this study, each metabolic reaction network encompasses approximately a hundred fluxes, representing various biochemical reactions occurring among different metabolites or within the same metabolite but across different cellular compartments. These fluxes comprehensively span the major cellular pathways, including glycolysis, the tricarboxylic acid (TCA) cycle, the pentose phosphate pathway (PPP), one-carbon metabolism (OCM), and glutamate/glutamine metabolism. To better emulate real cellular conditions, most metabolic networks are compartmentalized into three distinct regions: extracellular (e), cytosol (c), and mitochondria (m). Corresponding exchange fluxes facilitate the transfer of metabolites between these compartments. Each metabolite or reaction within the network is named by an uppercase abbreviation, followed by its compartment identifier. For instance, "GLC_c" signifies glucose in the cytosol, while "ICDH_m" designates isocitrate dehydrogenase in the mitochondria. Reversible fluxes are handled by regarding them as two independent fluxes, and the reverse reaction is labeled with the suffix "__R" (e.g., "LDH_c" and "LDH_c__R"). This naming convention ensures that all fluxes remain non-negative. In theory, flux values can be any non-negative quantity, but in practice, for the sake of simplicity and biological relevance, they are typically constrained within specific ranges defined by lower and upper bounds. A comprehensive table containing the full names and corresponding abbreviations of metabolites and fluxes is available in the file `model_metabolites_reactions_standard_name/standard_name.xlsx`.

A fundamental concept in flux analysis is the flux balance assumption, which posits that a cell will maintain a steady state concentration of each metabolite. Consequently, it is imposed as a constraint that, for the majority of metabolites, the sum of input fluxes equals the sum of output fluxes (as expressed in Eq. S1). There are exceptions, where specific metabolites are considered infinitely large pools acting as sources or sinks, such as CO2, extracellular 13C-labeled glucose, and lactate. The metabolites exempt from flux balance constraints are outlined in each analysis. Mathematically, this constraint can be represented as:

$$\sum_{\forall j} v_{ji} = \sum_{\forall k} v_{ik}, \text{ while } v_{ji} \text{ denotes the flux from metabolite } j \text{ to metabolite } i \qquad \text{(S1)}$$

After considering all these constraints, a feasible solution is in the form of a vector $\boldsymbol{v} = [v_1, v_2, \cdots, v_a]$, which contains the values of a total of $a$ fluxes that satisfy the flux balance constraints within the metabolic reaction network.

#### Mass isotopomer distribution calculation

The Mass Isotopomer Distribution (MID) for each metabolite is a vector obtained through mass spectrometry (MS), representing the ratios of various mass isotopologues resulting from isotope-labeling experiments. In this study, all labeling experiments exclusively employ the $^{13}$C isotope, making each mass isotopologue distinguishable by its count of $^{13}$C carbons. Within the MID vector, each element quantifies the proportion of the metabolite's mass distribution, ranging from $m + 0$ to $m + c$, where $m$ signifies the mass number of the raw metabolite, and $c$ denotes the total carbon count within that specific metabolite. Conceptually, the MID vector of a metabolite is shaped by and can be predicted from the weighted average of its upstream substrates, where the weights are determined by their respective generating fluxes. This relationship can be expressed as:

$$\widetilde{\boldsymbol{M}}_i = \frac{\sum_{\forall j} v_{ji} \boldsymbol{M}_{ji}}{\sum_{\forall j} v_{ji}}$$

$$\text{or } \left(\sum_{\forall j} v_{ji}\right) \widetilde{\boldsymbol{M}}_i = \sum_{\forall j} \left(v_{ji} \boldsymbol{M}_{ji}\right) \quad \text{(S2)}$$

In this equation, $\widetilde{\boldsymbol{M}}_i$ represents the predicted MID vector for metabolite $i$, $\boldsymbol{M}_{ji}$ stands for the MID of metabolite $i$ produced from substrate $j$, and $v_{ji}$ corresponds to the flux from $j$ to $i$.

Eq. S2 is applicable to most metabolites. However, certain metabolites, as mentioned in the "Flux model" section, are treated as infinitely large sinks or sources, such as $CO_2$ and extracellular metabolites. Given that any input or output flux for these metabolites is negligible compared to their pool size, the MID for these metabolites should be set to a fixed value. These metabolites are referred to as "input metabolites". Among these input metabolites, the MID for $^{13}$C-labeled substrates should match their labeling pattern. For instance, U-$^{13}$C-glucose exhibits an MID of $[0, 0, 0, 0, 0, 0, 1]$. In contrast, the MID of other unlabeled input metabolites with a carbon count of $k$ should follow a natural distribution, denoted as $\boldsymbol{M}_{\text{nat}}(k)$. This distribution is based on the binomial distribution principle, taking into account the natural abundance of $^{13}$C, denoted as $R_{13\text{C}}$ (approximately 1.109%). The formula for the natural distribution $\boldsymbol{M}_{\text{nat}}(k)$ is as follows:

$$M_{\text{nat},\, i}(k) = \binom{k}{i} R_{13\text{C}}^{i} \left(1 - R_{13\text{C}}^{i}\right)^{k-i},\ i = 0, 1, \ldots, k \quad \text{(S3)}$$

For example, the MID of unlabeled serine in the extracellular space (SER_e) is represented as $[0.9671, 0.03254, 3.649\text{e-}4, 1.3639\text{e-}6]$. As a result, based on Eq. S2, given a specific flux vector $\boldsymbol{v}$, the

MIDs of all metabolites can ultimately be expressed as linear equations involving the MIDs of these input metabolites. This is the reason they are referred to as "input metabolites".

The procedures described above represent a conceptual framework. In practical applications, simplification algorithms are often employed to streamline the computational intensity involved in large metabolic networks. One widely used computational framework for efficiently predicting MIDs of all metabolites is the Elementary Metabolite Units (EMU) algorithm[1]. The EMU algorithm divides the carbon chain of each metabolite into smaller units with different combinations of carbon atoms, referred to as EMUs. Notably, a complete metabolite can be viewed as an EMU that encompasses all carbon atoms. These EMUs are treated as independent metabolites. Their substrates are traced, their MIDs are computed based on Eq. S2, and the results are recorded. These calculated and recorded MIDs can then be directly reused when calculating other EMUs that stem from the same combination of carbon atoms, eliminating the need for redundant computations. Tracing the upstream substrates of EMUs relies on carbon mapping equations, which document the transfer of carbon from substrate to product in each metabolic flux within the model.

With a complete list of reactions and their corresponding carbon mapping equations, the EMU algorithm generates a set of linear equations ordered by the carbon number of EMUs. These equations enable the determination of MIDs for all EMUs (and consequently, metabolites) denoted as $\widetilde{\boldsymbol{M}}_i$, given a predefined flux vector $\boldsymbol{v}$, the MIDs of input metabolites $\boldsymbol{M}_{\text{input},j}$, and previously solved MIDs of smaller EMUs, $\widetilde{\boldsymbol{M}}_j$. This relationship can be expressed as:

$$\boldsymbol{A}(\boldsymbol{v})\big[\widetilde{\boldsymbol{M}}_i\big] = \boldsymbol{B}(\boldsymbol{v})\big[\boldsymbol{M}_{\text{input},j} \text{ or } \widetilde{\boldsymbol{M}}_j\big] \tag{S4}$$

Here, $\boldsymbol{A}(\boldsymbol{v})$ and $\boldsymbol{B}(\boldsymbol{v})$ represent matrices determined by the flux vector $\boldsymbol{v}$. Eq. S4 can be conceptualized as stacking multiple instances of Eq. S2 vertically and organizing these linear relationships into matrix form. $\boldsymbol{A}(\boldsymbol{v})$ and $\boldsymbol{B}(\boldsymbol{v})$ are essentially matrix representations of $v_{ji}$ as found in Eq. S2.

In this study, the EMU algorithm is implemented in a computationally efficient manner. Since the MIDs of input metabolites remain fixed within a single experiment, the EMU-generated linear equations (Eq. S4) are structured to accommodate flux values as variables while treating other parameters as constants. The locations of flux in $\boldsymbol{A}(\boldsymbol{v})$ and $\boldsymbol{B}(\boldsymbol{v})$ are left blank. After filling $\boldsymbol{A}(\boldsymbol{v})$ and $\boldsymbol{B}(\boldsymbol{v})$ in Eq. S4 with a given flux vector, the MIDs of all metabolites in the metabolic network can be uniquely and efficiently predicted. Therefore, in the subsequent paragraphs, each predicted $\widetilde{\boldsymbol{M}}_i$ of metabolite $i$ can be considered a function of the flux vector $\boldsymbol{v}$, marked as $\widetilde{\boldsymbol{M}}_i(\boldsymbol{v})$.

#### Input metabolites in experimental analyses

As mentioned above, input metabolites typically include external nutrients absorbed by cells or waste products secreted out, such as extracellular glucose, lactate, glutamine, and CO2. MIDs of these metabolites are considered invariant, irrespective of the flux magnitude entering them.

Lactate, among these, presents a unique case. In glucose tracing experiments, the actual MID of extracellular lactate is influenced by labeled intracellular lactate. However, our model assumes that lactate is unidirectionally output, meaning the 13C labeling of extracellular lactate has minimal effect on our results. While this may slightly underestimate lactate reuse by tissues, any reutilization shares identical MID values with cellular lactate. Hence, this effect is accounted for in cellular lactate MID fitting, and the lactate output flux in our model can be viewed as the net lactate release rate from cancer cells.

The 13C labeling in other input metabolites, including glutamine, aspartate, serine and glycine, ultimately originates from 13C labeled glucose. However, due to the relatively low labeling ratio of glucose, the impact of this labeling on these additional metabolites is negligible and can be safely disregarded in our analysis.

Our data sources sometimes include plasma MIDs of these metabolites, but we have chosen not to incorporate them into our analysis for two main reasons. First, all measurements in isotope tracing are susceptible to noise, introduced during data acquisition stages such as sample collection, preparation, and measurement. If MIDs in plasma were treated as input data to predict other MIDs for fitting, their inherent noise would propagate to the MIDs of other metabolites, potentially compromising the accuracy of the model.

Secondly, the EMU algorithm used for predicting MIDs from flux requires the knowledge of the labeling ratio of each carbon atom for input metabolites. However, experimental measurements can only provide MIDs for complete metabolites rather than the individual carbon atoms within them. As a result, these data cannot be directly utilized in the EMU algorithm.

#### MID mixing and mapping

The predicted MIDs of metabolites within the model must be aligned with experimental datasets for comparison. When the MID of a metabolite can be measured in the dataset and can be uniquely linked to a predicted MID in the flux model, a direct comparison between the experimental and predicted MIDs can be executed to calculate the final loss for that specific metabolite. In real isotope tracing data, it is common that only a subset of predicted MIDs, often fewer than 30, can be mapped to experimental ones that contributes to the final loss function.

Experimental MIDs of metabolites are usually measured at the whole-cell level. However, in compartmental metabolic models, some metabolites may exist in multiple cellular compartments, each having a different predicted MID. For instance, metabolites like α-ketoglutarate, aspartate, glutamate, and malate can be found in both the cytosol and mitochondria, but experimental measurements are often taken as whole-cell averages in most datasets. Apart from compartmental metabolites, certain isomers cannot be distinguished by mass spectrometry, such as citrate/isocitrate and 2-phosphoglycerate/3-phosphoglycerate, resulting in their MIDs being measured at an average level.

To account for these scenarios, an additional set of independent pool size variables is introduced for each metabolite. The mixed MID is then computed as an average weighted by these pool sizes, defined as follows:

$$\widetilde{\boldsymbol{M}}_i(\boldsymbol{v},\boldsymbol{p}) = \frac{\sum_{\forall k} p_k \widetilde{\boldsymbol{M}}_{i,k}(\boldsymbol{v})}{\sum_{\forall k} p_k} \tag{S5}$$

Here, $\widetilde{\boldsymbol{M}}_i(\boldsymbol{v},\boldsymbol{p})$ represents the mixed MID vector for metabolite $i$, $\widetilde{\boldsymbol{M}}_{i,k}(\boldsymbol{v})$ stands for the predicted MID of any elemental metabolite $k$ that contributes to the mixture in metabolite $i$, and $p_k$ signifies the pool size variable for metabolite $k$.

The calculation of the mixed distribution (Eq. S5) closely resembles the calculation of the predicted MID (Eq. S4). These two calculations are designed, constrained, and carried out using similar strategies. Similar to flux variables represented by $\boldsymbol{v}$, pool size variables $p_k$ are determined through optimization. Since $p_k$ denotes the pool size of metabolite $k$ and remains the same in multiple mixing equations that metabolite $k$ participates in (commonly encountered when simultaneously fitting multiple experimental datasets with the same flux vector), a mixed vector $\boldsymbol{p}$ can also be defined, comprising values such as

$[p_1, p_2, \cdots, p_l]$. Like fluxes, theoretically, $p_k$ could assume any non-negative value, but in practice, they are typically constrained within specified lower and upper bounds. Moreover, $\sum_{\forall k} p_k$ is fixed to a constant value across all mixing equations. The notation $\widetilde{\boldsymbol{M}}_i(\boldsymbol{v}, \boldsymbol{p})$ is used to signify that the mixed predicted distribution is a function of both the flux vector $\boldsymbol{v}$ and the mixture vector $\boldsymbol{p}$.

#### Natural isotope abundance in experimental MID data

The natural abundance of $^{13}$C is approximately 1.109%. Consequently, there is a non-zero $^{13}$C labeling ratio even in unlabeled metabolites. Many contemporary studies address this issue by correcting for the natural abundance of 13C in the experimental MID prior to any subsequent analyses. In this protocol, the raw measured MID distribution, denoted as $\boldsymbol{M}_r$, is adjusted by calculating the natural distribution of the same metabolite, $\boldsymbol{M}_n$, using Eq. S3. The first element of $\boldsymbol{M}_n$ is then replaced by 0 to create the corrected distribution, denoted as $\widehat{\boldsymbol{M}}_n$, as shown in the following equation:

$$\widehat{M}_{n,i} = \begin{cases} 0 & i = 0 \\ M_{n,i} = \binom{k}{i} R_{13\text{C}}^{i} \left(1 - R_{13\text{C}}^{i}\right)^{k-i} & i = 1,2,\dots,k \end{cases} \tag{S6}$$

The correction function, Cor, and the corrected MID distribution, $\boldsymbol{M}_c$, are defined by normalizing the direct subtraction between $\boldsymbol{M}_r$ and $\widehat{\boldsymbol{M}}_n$, as described in the following equation:

$$\boldsymbol{M}_c = \mathrm{Cor}(\boldsymbol{M}_r) = \frac{\boldsymbol{M}_r - \widehat{\boldsymbol{M}}_n}{\sum(\boldsymbol{M}_r - \widehat{\boldsymbol{M}}_n)} = \frac{1}{M_{n,0}}\left(\boldsymbol{M}_r - \widehat{\boldsymbol{M}}_n\right) \tag{S7}$$

Here, $k$ represents the carbon number of the current metabolite, and $M_{n,0}$ denotes the raw 0-th element of the natural distribution $\boldsymbol{M}_n$. It's important to note that the correction function Cor is a linear transformation that converts the naturally distributed MID $\boldsymbol{M}_n$ to a fully unlabeled MID $[1,0,\dots,0]$.

In real metabolic networks, considering a simple scenario where a metabolite $Z$ has two substrates, including metabolite $X$ with MID $\boldsymbol{M}_X$ and flux $v_1$, and metabolite $Y$ with MID $\boldsymbol{M}_Y$ and flux $v_2$, the raw measured MID of metabolite $Z$, referred to as $\boldsymbol{M}_Z$, can be expressed by the following equation:

$$\boldsymbol{M}_Z = \frac{v_1}{v_1 + v_2}\boldsymbol{M}_X + \frac{v_2}{v_1 + v_2}\boldsymbol{M}_Y \tag{S8}$$

Since both this equation and the correction function Cor are linear functions, it becomes evident that:

$$\mathrm{Cor}(\boldsymbol{M}_Z) = \frac{v_1}{v_1 + v_2}\mathrm{Cor}(\boldsymbol{M}_X) + \frac{v_2}{v_1 + v_2}\mathrm{Cor}(\boldsymbol{M}_Y) \tag{S9}$$

Hence, if all MIDs can be corrected using Eq. S7, the corrected MIDs are entirely equivalent to the uncorrected raw MIDs when calculating a flux model.

However, these assumptions underlying the correction function Cor in Eq. S7 may not always hold. First, it assumes that the MID of all metabolites includes the natural abundance of 13C. However, this assumption doesn't hold for labeled substrates, such as U-13C-glucose, which are synthetically produced and guaranteed to be fully labeled. Secondly, this correction function requires that all isotopologues in the measured MID, $\boldsymbol{M}_r$, are greater than the corresponding values in $\widehat{\boldsymbol{M}}_n$. Otherwise, negative values may arise in the corrected MID $\boldsymbol{M}_c$. In practical applications, negative values tend to emerge in metabolites that are in close proximity to labeled input metabolites in the flux model, such as glucose 6-phosphate, since their measured MIDs are typically fully labeled. Negative values can also occur in metabolites with a greater number of carbon atoms, like fatty acids, where the $\widehat{\boldsymbol{M}}_n$ values for m+1/2 isotopologues are relatively larger. Negative values can disrupt the normalization property of the MID and render the calculation of certain loss functions, such as Kullback–Leibler divergence, infeasible (as discussed in the "Loss function calculation" section below). Therefore, for enhanced numerical stability, in this study, as noted in the "Mass isotopomer distribution calculation" section, we account for the natural abundance of $^{13}$C in all MID calculations, and all unlabeled MIDs are set to the natural distribution as defined in Eq. S3.

Some of the published data used in this study have undergone correction, but neither the raw data nor detailed correction protocols have been provided in their respective papers. Consequently, we assume that their correction process is based on Eq. S7 and employ an inverse transformation to reconstruct $\boldsymbol{M}_r$. Following the inverse transformation, any elements in $\boldsymbol{M}_r$ that are negative or exceed 1 are clipped into the $[0,1]$ range and then normalized once more. The resulting processed MID data are subsequently utilized in the subsequent analyses.

In addition to $^{13}$C, the measurement of MID can be affected by the presence of rare natural isotopes of other elements, such as nitrogen ($^{15}$N), sulfur ($^{33}$S, $^{34}$S), hydrogen ($^{2}$H), and oxygen (18O). However, the impact of these rare natural isotopes on MID measurements depends on both their abundance and their occurrence in metabolites. It is worth noting that our flux model excludes metabolites that contain sulfur (such as methionine, cysteine, or biotin). Regarding nitrogen atoms, our model does not encompass basic amino acids or bases, ensuring that metabolites in our model contain no more than one nitrogen atom. Consequently, the presence of the 0.4% $^{15}$N abundance has an insignificant influence on the final MIDs of metabolites. As for hydrogen, any biological molecule with a carbon backbone cannot have four times as many hydrogen atoms as carbon atoms. Given its 0.01% abundance, the impact of $^{2}$H on final MIDs can be confidently disregarded. Similarly, the abundance of the $^{18}$O isotope is limited (0.2%), and the number of oxygen atoms in metabolites is hardly significantly greater than the number of carbon atoms. Therefore, for the majority of metabolites containing fewer than six carbon atoms, the contribution of the $^{18}O$ isotope to the final MIDs is approximately 1%, still falling below the typical error rate of MID data.

#### Loss function calculation

The primary objective of MFA is to minimize the disparity between the predicted and experimental MID data. This discrepancy between the predicted (mixed) MID $\widetilde{\boldsymbol{M}}_i(\boldsymbol{v},\boldsymbol{p})$ and the experimentally observed MID $\boldsymbol{M}_i$ for a given metabolite $i$ can be effectively quantified using the Kullback–Leibler divergence, $D_{\mathrm{KL}}$[2]. This measure of disparity is commonly referred to as the loss function $L_i(\boldsymbol{v},\boldsymbol{p})$:

$$L_i(\boldsymbol{v},\boldsymbol{p}) = D_{\mathrm{KL}}\left(\widetilde{\boldsymbol{M}}_i(\boldsymbol{v},\boldsymbol{p}) \parallel \boldsymbol{M}_i\right) = \sum_j \left(M_{i,j} + \varepsilon_{\log}\right) \log \frac{\widetilde{M}_{i,j}(\boldsymbol{v},\boldsymbol{p}) + \varepsilon_{\log}}{M_{i,j} + \varepsilon_{\log}} \tag{S10}$$

In this equation, $M_{i,j}$ and $\widetilde{M}_{i,j}$ represent the $j$-th elements in the vectors $\boldsymbol{M}_i$ and $\widetilde{\boldsymbol{M}}_i(\boldsymbol{v},\boldsymbol{p})$, respectively. The term $\varepsilon_{\log}$ is a small value added to ensure numerical stability. It's important to note that this loss function assumes that the experimental MID $\boldsymbol{M}_i$ is positive for all isotopologues, which cannot always be guaranteed when natural abundance is directly corrected in experimental MID.

The overall total loss function, denoted as $L_{\mathrm{total}}$, is the summation of the loss functions for all mapped metabolites, expressed as:

$$L_{\mathrm{total}}(\boldsymbol{v},\boldsymbol{p}) = \sum_{\forall i} L_i(\boldsymbol{v},\boldsymbol{p}) \tag{S11}$$

In practical applications, there are typically numerous mapped metabolites. It is evident that $L_{\mathrm{total}}$ is uniquely determined by the flux vector $\boldsymbol{v}$ and mix vector $\boldsymbol{p}$. The optimization algorithm leverages $L_{\mathrm{total}}$ to update $\boldsymbol{v}$ and $\boldsymbol{p}$, ultimately seeking the optimal solutions.

#### Constraints and definition of MFA problem

As previously mentioned, the flux vector $\boldsymbol{v}$ must adhere to flux balance constraints, which necessitate that the total input flux equals the total output flux for most metabolites, and must also satisfy lower and upper bounds. Similarly, the mix vector $\boldsymbol{p}$ is subject to summation constraints, requiring that $\sum_{\forall k} p_k$ equals a predetermined value, with lower and upper bounds to be respected. All these constraints are linear in nature.

Additionally, there are several supplementary linear constraints that the solution must adhere to. Firstly, since the flux model is invariant when all fluxes are multiplied by a constant, any solution obtained through MFA represents a relative value. Consequently, in MFA analyses, a specific flux is typically set as a fixed value during analysis. This enables the calculation of other fluxes relative to this fixed value. In the analyses conducted in this study, one or more input fluxes, such as those associated with glucose, glutamine, and aspartate, are often set as fixed fluxes. The selection of these fixed fluxes can influence the precision and accuracy of the solution.

Secondly, under biological conditions, different fluxes can exhibit significantly different dynamic ranges. For example, primary glycolytic fluxes often fall within the category of the largest fluxes, particularly in cancer cells, where they can be approximately 100 times higher than other metabolic pathways, such as fatty acid synthesis. To address this, it is important to establish specific bounds for known smaller fluxes to reduce the solution space and prevent the solution from becoming trapped in incorrect local optima. In addition to these scenarios, supplementary linear constraints may also be introduced, such as flux values and linear relationships determined by thermodynamics or data from other experiments.

Flux ranges, often referred to as bounds, must be established for all fluxes prior to the optimization process. Typically, the flux range for normal fluxes falls within the interval $[1{,}1000]$ in most optimization cases. For mixing fluxes, as previously mentioned, they have their own predefined flux range, which is usually set to $[5{,}95]$ to ensure that the fitted values for mixing fluxes are similar with the percentage number. Additional linear flux constraints and adjustments to flux ranges are common configuration parameters that may impact the solutions obtained during the optimization process, and as a result, these parameters are subject to perturbation in parameter sensitivity analyses.

As both the flux vector $\boldsymbol{v}$ and mix vector $\boldsymbol{p}$ are mathematically equivalent within this problem, for the sake of simplicity, we will use the composite vector $\boldsymbol{v}$ to represent the combination $[\boldsymbol{v}, \boldsymbol{p}]$. Taking into account the loss function and all constraints, the problem of MFA can be formally defined as a linear constrained nonlinear optimization problem, described by the following set of equations:

$$\begin{gathered} \min_{\boldsymbol{v}} L_{\text{total}}(\boldsymbol{v}) \\ \text{s.t. } \boldsymbol{A} \cdot \boldsymbol{v}^T = \boldsymbol{b} \\ \boldsymbol{C} \cdot \boldsymbol{v}^T \leq \boldsymbol{d} \\ \boldsymbol{0} \leq \boldsymbol{v}_{\min} \leq \boldsymbol{v} \leq \boldsymbol{v}_{\max} \end{gathered} \tag{S12}$$

In this formulation, $\boldsymbol{A}$ and $\boldsymbol{b}$ represent the equality linear constraints, while $\boldsymbol{C}$ and $\boldsymbol{d}$ denote the inequality linear constraints. The values $\boldsymbol{v}_{\min}$ and $\boldsymbol{v}_{\max}$ establish the lower and upper bounds for the composite vector $\boldsymbol{v}$. It's important to recognize that the loss function $L_{\text{total}}(\boldsymbol{v})$ is inherently a non-convex nonlinear function, making it challenging to find the global optimal solution. Consequently, a precise and consistent optimization algorithm remains one of the most complex and demanding aspects of the MFA problem.

#### Fixed fluxes in experimental analyses

As mentioned above, a crucial aspect of MFA is the proportional relationship among all fluxes in the results. For meaningful comparisons across different samples, it's essential to have a consistent standard or fixed flux that is comparable in all samples. In glucose tracing experiments, which form the basis of our simulated data analyses, the glucose input flux is often used as this fixed standard.

However, in cancer tissues, glucose uptake can vary significantly due to the high energy demands of cancer cells, diminishing its reliability as a standard for flux comparison. Similarly, in cultured cells, limited glucose availability can naturally reduce glucose uptake flux. Therefore, in analyzing experimental datasets in our study, we employ different fixed fluxes. Our analysis suggests that a constant aspartate intake flux can stabilize many smaller fluxes, reducing their variability across samples. The specific value of this aspartate uptake flux is less important than its consistent presence across different samples. Consequently, we have chosen to set aspartate uptake flux as a fixed value in our experimental dataset analysis. This approach ensures a stable reference point for comparisons.

#### Feasible solution generator

Obtaining a feasible solution, which is a vector $\boldsymbol{v}$ that complies with all the constraints outlined in Eq. S12, is a critical step in many nonlinear optimization algorithms. This study involves comparing random feasible solutions with optimized results, which necessitates the efficient generation of thousands of feasible solutions in a high-dimensional space. However, it's crucial to understand that the solution space for metabolic networks can be extraordinarily vast. For example, considering the network depicted in Figure 1C, there are approximately 100 flux variables but only about 50 linear constraints, including flux balance constraints and fixed value constraints. The degree of freedom of this problem is therefore around 50, implying that the size of the solution space is approximately $10^{150}$ (considering a flux range from 1 to 1000). Even though all constraints are linear, and the feasible region is convex, the time complexity of a brute-force algorithm seeking a solution that satisfies all constraints is exponential. The running time of such an algorithm would be impractical for the scale of this study.

A more efficient approach for generating feasible solutions within a convex linear space is the application of linear programming (LP) with a random objective function. This involves solving the following problem:

$$\min_{\boldsymbol{v}} \boldsymbol{r} \cdot \boldsymbol{v}^T$$
$$\text{s.t. } \boldsymbol{A} \cdot \boldsymbol{v}^T = \boldsymbol{b}$$
$$\boldsymbol{C} \cdot \boldsymbol{v}^T \leq \boldsymbol{d}$$
$$\boldsymbol{v}_{\min} \leq \boldsymbol{v} \leq \boldsymbol{v}_{\max} \tag{S13}$$

Here, $\boldsymbol{r}$ represents a non-zero random vector with the same dimension as the composite variable vector $\boldsymbol{v}$ (i.e., $[\boldsymbol{v}, \boldsymbol{p}]$). The linear programming problem described in Eq. S13 can be effectively solved using the simplex algorithm implemented in the SciPy package[3,4]. It's important to note that LP-based methods have a specific property – they tend to generate feasible solutions that are situated near the corners of the solution space. While this approach is efficient, it may not be ideal for producing roughly uniformly distributed random solutions, and this characteristic could potentially hinder subsequent optimization algorithms.

A more effective strategy involves using the solutions obtained from Eq. S13 as seeds and implementing an iterative algorithm for generating additional feasible solutions from these seeds. One widely used approach for this purpose is the hit-and-run (HR) algorithm[5]. The fundamental concept of the HR algorithm is as follows: starting from a feasible solution, a point moves in a random direction by a random distance. If the new position adheres to the constraints, it is stored as a new feasible solution. There are various derivative forms of the hit-and-run algorithm suitable for different types of problems. In this study, we adapt an HR-derived algorithm previously developed and known as optGpSampler[6]. We implement a sampling algorithm tailored to the problem defined in Eq. S12. The detailed procedures are as follows:

Algorithm S1:

1. Initialize a pool of feasible solutions $V = \{\boldsymbol{v}'_0\}$ generated by solving the LP problem (Eq. S13). The pool size $n$ should be greater than the final required sample size.

2. Warm-up steps:
    a. Randomly select multiple feasible solutions in $V$ and calculate their mean vector $\boldsymbol{v}_{\text{start}}$ as the initial point of the sampler. It's evident that $\boldsymbol{v}_{\text{start}}$ also adheres to all constraints in Eq. S12. Set $\boldsymbol{v} = \boldsymbol{v}_{\text{start}}$.
    b. Choose a uniformly random direction and map it to the null space of matrix $\boldsymbol{A}$ in Eq. S12. This derived direction is referred to as $\boldsymbol{d_A}$. By definition of the null space, $\boldsymbol{A} \cdot \boldsymbol{d_A} = \boldsymbol{0}$.
    c. Starting from $\boldsymbol{v}$, search along $\boldsymbol{d_A}$ to find the maximal distance $\alpha_{\max}$ such that $\boldsymbol{v}_{\text{new, max}} = \boldsymbol{v} + \alpha_{\max} \cdot \boldsymbol{d_A}$ still adheres to the inequality constraints $\boldsymbol{C} \cdot \boldsymbol{v}_{\text{new, max}}{}^{T} \leq \boldsymbol{d}$ and the bounds $\boldsymbol{v}_{\min} \leq \boldsymbol{v}_{\text{new, max}} \leq \boldsymbol{v}_{\max}$.
    d. Draw a uniformly random number $\alpha$ from the interval $(0, \alpha_{\max})$. Calculate the updated location $\boldsymbol{v}_{\text{new}} = \boldsymbol{v}_{\text{start}} + \alpha \cdot \boldsymbol{d_A}$.
    e. Set $\boldsymbol{v} = \boldsymbol{v}_{\text{new}}$ and return to step b. After every $t$ repetitions, replace one random element in $V$ with the current $\boldsymbol{v}$ (referred to as the thinning number). End after $n$ replacements.
3. Formal sampling steps:
    a. Calculate the mean vector $\boldsymbol{v}_{\text{mean}}$ of all vectors in $V$. It's evident that $\boldsymbol{v}_{\text{mean}}$ also adheres to all constraints in Eq. S12.
    b. Randomly select a previous feasible solution $\boldsymbol{v}_{\text{random}}$ from $V$ and calculate the direction vector $\boldsymbol{d} = \boldsymbol{v}_{\text{random}} - \boldsymbol{v}_{\text{mean}}$. Because $\boldsymbol{v}_{\text{random}}$ and $\boldsymbol{v}_{\text{mean}}$ are two feasible solutions, $\boldsymbol{A} \cdot \boldsymbol{d} = \boldsymbol{0}$.
    c. Similar to step 2c, replace $\boldsymbol{d_A}$ with $\boldsymbol{d}$ and calculate $\alpha_{\max}$.
    d. Similar to step 2d, draw a random $\alpha$ and calculate $\boldsymbol{v}_{\text{new}}$.
    e. Similar to step 2e, update $\boldsymbol{v}$ and return to step b. Replace an element in $V$ with the thinning number $t$ and update $\boldsymbol{v}_{\text{mean}}$ after replacement. End after more than $n$ replacements.
4. Randomly select the final required sample size from the set $V$.

In this study, all initial points for optimization are generated using Alg. S1. Random feasible fluxes utilized in Fig. 1 and S1 are also generated by this algorithm.

The vast solution space, as mentioned above, means that only thousands of samples may not necessarily be widely dispersed and may not adequately represent the entire solution space. This might be the main reason that random feasible fluxes are usually distantly distributed from the optimized solutions (Figure S1B)

#### Optimization

Eq. S12 defines a non-convex nonlinear optimization problem with linear constraints. The quest for a global optimum in such problems is known to be at least NP-hard, which implies that there is no algorithm that can guarantee the discovery of the global optimum. However, there is a silver lining in the context of MFA. The loss function employed in MFA exhibits certain favorable characteristics for optimization purposes: it is continuous, infinitely differentiable, and characterized by a lack of steep variations across its entire defined domain. These properties enhance the likelihood of finding satisfactory local-optimum solutions when employing a well-suited optimization algorithm.

In this study, we employ the Sequential Least Squares Programming (SLSQP) algorithm, which is integrated into the scipy.optimize.minimize function within the SciPy package[7]. Designed for the optimization of non-convex nonlinear functions with nonlinear constraints, the algorithm tackles problems defined as follows:

$$\begin{aligned} &\min_{\boldsymbol{v}} f(\boldsymbol{v}) \\ \text{s.t. } &\boldsymbol{b}(\boldsymbol{v}) \geq \boldsymbol{0} \\ &\boldsymbol{c}(\boldsymbol{v}) = \boldsymbol{0} \end{aligned} \tag{S14}$$

Here, $f$, $\boldsymbol{b}$ and $\boldsymbol{c}$ represent various functions. Such problems are typically addressed using Sequential Quadratic Programming (SQP), where the Lagrangian is defined as:

$$\mathcal{L}(\boldsymbol{v}, \boldsymbol{\lambda}, \boldsymbol{\sigma}) = f(\boldsymbol{v}) - \boldsymbol{\lambda}^T \boldsymbol{b}(\boldsymbol{v}) - \boldsymbol{\sigma}^T \boldsymbol{c}(\boldsymbol{v})$$

In this context, $\boldsymbol{\lambda}$ and $\boldsymbol{\sigma}$ are Lagrangian multipliers. The SLSQP algorithm commences from a feasible solution $\boldsymbol{v}_0$. In each iteration $k$, when at point $\boldsymbol{v}_k$, it computes the search direction $\boldsymbol{d}_k$ by solving the subproblem:

$$\begin{aligned} &\min_{\boldsymbol{d}_k} \nabla f(\boldsymbol{v}_k)^T \boldsymbol{d}_k + \tfrac{1}{2} \boldsymbol{d}_k^T \nabla^2 \mathcal{L}(\boldsymbol{v}_k, \boldsymbol{\lambda}_k, \boldsymbol{\sigma}_k) \boldsymbol{d}_k \\ \text{s.t. } &\boldsymbol{b}(\boldsymbol{v}_k) + \nabla \boldsymbol{b}(\boldsymbol{v}_k)^T \boldsymbol{d}_k \geq \boldsymbol{0} \\ &\boldsymbol{c}(\boldsymbol{v}_k) + \nabla \boldsymbol{c}(\boldsymbol{v}_k)^T \boldsymbol{d}_k = \boldsymbol{0} \end{aligned} \tag{S15}$$

In practice, the objective function $\min_{\boldsymbol{d}_k} \nabla f(\boldsymbol{v}_k)^T \boldsymbol{d}_k + \frac{1}{2} \boldsymbol{d}_k^T \nabla^2 \mathcal{L}(\boldsymbol{v}_k, \boldsymbol{\lambda}_k, \boldsymbol{\sigma}_k) \boldsymbol{d}_k$ can be reformulated into a least squares form, $\min_{\boldsymbol{d}_k} \frac{1}{2} \|\boldsymbol{G} \cdot \boldsymbol{d}_k - \boldsymbol{F}\|$. This transformation gradually converts the problem into a non-negative least square form, making it feasible to solve for $\boldsymbol{d}_k$[7]. The step size $\alpha_k$ at iteration $k$ is determined using a special metric called a merit function[8]. Finally, the updated point is computed as:

$$\boldsymbol{v}_{k+1} = \boldsymbol{v}_k + \alpha_k \, \boldsymbol{d}_k \quad \text{(S16)}$$

Subsequent iterations continue until $\|\boldsymbol{d}_k\| < \varepsilon$ or $k \geq k_{\max}$, which we have set as $k_{\max} = 500$ in our analysis. The final $\boldsymbol{v}_k$ is returned as the optimized solution.

In practical usage of the SLSQP algorithm, the problem defined in Eq. S12 is transformed into the standard form given by Eq. S14. Constraints in Eq. S12 that are less than or equal to are converted into greater-than-or-equal-to constraints in Eq. S14 by multiplying both sides by $-1$. The SLSQP function also accommodates variable bounds and automatically converts them into inequality constraints.

Much like other nonlinear optimization algorithms, SLSQP requires an initial feasible solution to start optimization and can only guarantee a local optimum from this initial point. Notably, SLSQP does not depend on any random factors. Consequently, the choice of an initial feasible solution becomes critically important.

Due to its detailed approach in each iteration, SLSQP is adept at capturing the intricacies of the solution space, often resulting in high-quality solutions. However, this precision comes at the cost of longer execution times and increased computational resource demands. Moreover, the algorithm's implementation in Fortran code may limit its ability to incorporate the latest advancements in computation.

#### Flux solution processing and embedding

In our model, numerous reversible fluxes exist. However, our primary focus is often on the net flux within each reversible pair rather than the absolute values of the individual reactions. Consequently, when we compare the model's fluxes with known data or visualize the final solutions, an additional step is introduced. The net flux is calculated by taking the difference between the two reactions within a reversible pair. The transformation $N(\boldsymbol{v})$ that maps the raw flux vector $\boldsymbol{v}$ to the net flux vector $\boldsymbol{v}_{\text{net}}$ can be expressed as a linear operation:

$$\boldsymbol{v}_{\text{net}} = N(\boldsymbol{v}) = \boldsymbol{P} \cdot \boldsymbol{v}$$
$$v_{\text{net},j} = \begin{cases} v_i - v_{i+1} \text{ or } v_{i+1} - v_i & \text{flux } i \text{ and } i+1 \text{ are pair of reversible fluxes} \\ v_i & \text{flux } i \text{ is unidirectional} \end{cases} \quad \text{(S17)}$$

The dimension of $\boldsymbol{v}_{\text{net}}$ is typically smaller than that of $\boldsymbol{v}$, which means that the transformed coordinate $j$ in $\boldsymbol{v}_{\text{net}}$ for flux $i$ (and $i+1$) in $\boldsymbol{v}$ is typically smaller than $i$. The direction (i.e., whether it is $v_i - v_{i+1}$ or $v_{i+1} - v_i$) is determined to ensure that the net flux is positive in as many solutions as possible. The matrix $\boldsymbol{P}$ represents the linear transformation matrix with elements $1$ and $-1$ at the locations corresponding to the reversible flux pairs, $1$ for all unidirectional fluxes, and $0$ elsewhere.

In the section titled "Assessment of current automated analysis method in labeling experiments" within the main text, we combine random and optimized solutions, embedding them through principal component analysis (PCA). This is done using the sklearn.decomposition.PCA function from the scikit-learn Python package. We generate a 2D scatter plot with two output components.

#### Raw and net Euclidean distance

The calculation of the Euclidean distance between two flux vectors $\boldsymbol{v}_1$ and $\boldsymbol{v}_2$, referred as $D(\boldsymbol{v}_1, \boldsymbol{v}_2)$, is calculated by the L2-norm of the differential vector:

$$D(\boldsymbol{v}_1, \boldsymbol{v}_2) = \|\boldsymbol{v}_1 - \boldsymbol{v}_2\|_2 \tag{S18}$$

When the Euclidean distance is directly computed from the raw flux vector, it is termed as the raw Euclidean distance. Conversely, the Euclidean distance between net flux vectors, which are derived from the corresponding raw flux vectors using Eq. S17, is referred to as the net Euclidean distance. Since researchers typically prioritize the accuracy of net fluxes in MFA solutions, the net Euclidean distances, which better assess these differences, are emphasized in the following analyses.

In the visual representation of the results obtained from the HCT-116 dataset, both the raw and net Euclidean distances are showcased. These distances are calculated between the best solution with the minimal loss value and other best or random solutions, as presented in Figure 1H. Additionally, the detailed difference between these net fluxes is plotted in Figure S1C for further examination.

#### Simulated data generation

The predefined flux vector $\boldsymbol{v}'$, represents an optimized solution that originates from an initial random flux. This optimization process is informed by previously published MID data obtained from cultured cancer cells, specifically HCT116. A more detailed configuration of this optimization can be found in the section titled "MFA for HCT116 Cultured Cancer Cell Line" in the second part below.

The MID data is then predicted using this predefined flux vector $\boldsymbol{v}'$, as well as the same input metabolites used in the HCT116 dataset. The predicted MIDs of all metabolites, without considering any mixing variables, are harnessed to create a simulated MID dataset, representing all-available simulated MID data. Additionally, the predicted MIDs of metabolites that match the ones found in the HCT116 dataset and incorporate mixing variables are used to simulate experimentally-available MID data.

In our subsequent comparisons, we also calculate the net flux vector $\boldsymbol{v}'_{\text{net}}$ for $\boldsymbol{v}'$ using Eq. S17. For instance, if LDH_c has a flux value of 50 and LDH_c__R is at 20 in $\boldsymbol{v}'$, the direction of the net LDH flux is defined as LDH_c − LDH_c__R, resulting in a value of 30 for this particular flux in $\boldsymbol{v}'$.

To validate our algorithm's capability to identify predefined flux vectors across the entire solution space, we generated 30 predefined solutions that are geometrically distant from each other. Each pair of these solutions has a mutual Euclidean distance in net fluxes exceeding 500, ensuring their distinctiveness. These predefined flux vectors were then subjected to the same protocol analysis as outlined in the subsequent section. Their performance metrics, defined by loss and net Euclidean distance, are calculated based on $\mathbb{L}_{\text{sel}}(20000, 100, 1)$, $\mathbb{L}_{\text{ave}}(20000, 100, 1)$, $\mathbb{L}_{\text{re-op}}(20000, 100, 1)$, $\mathbb{D}_{\text{sel}}(20000, 100, 1)$, $\mathbb{D}_{\text{ave}}(20000, 100, 1)$ and $\mathbb{D}_{\text{re-op}}(20000, 100, 1)$ (these definitions can be found in the subsequent section) and displayed in Figure 2C, 2G.

#### MFA protocol analysis

In Figure 2B, we initiate the process by generating $n$ initial random flux vectors through Alg. S1. Independent optimizations are then conducted, from these initial flux solutions, through the SLSQP algorithm, yielding $n$ optimized results. Among these optimized results, we select the $m$ solutions with the minimal loss values. These selected solutions and their loss values, obtained under the current parameters $(m, n)$, is referred as $V_{\text{sel}}(m, n) = \{\boldsymbol{v}_{\text{sel}, p} | p = 1, \dots, m\}$ and $L_{\text{sel}}(m, n) = \{L_{\text{sel}, p} | p = 1, \dots, m\}$. Following this, we calculate the average flux vector, referred to as $\boldsymbol{v}_{\text{ave}}(m, n)$, from these $m$ selected solutions, and evaluate its loss value $L_{\text{ave}}(m, n)$. We also incorporate an additional optimization step beginning from $\boldsymbol{v}_{\text{ave}}(m, n)$, leading to what we term as the re-optimized solutions,$\boldsymbol{v}_{\text{re-op}}(m, n)$.

This entire optimization-averaging process is repeated $k = 30$ times. We combine selected solutions and their loss values in all repetitions as $\mathbb{V}_{\text{sel}}(m,n,k) = \{\cup V_{\text{sel},\,i}(m,n)\,|i = 1,\dots,k\}$ and $\mathbb{L}_{\text{sel}}(m,n,k) = \{\cup L_{\text{sel},\,i}(m,n)\,|i = 1,\dots,k\}$. At the same time, we also collect all averaged solutions and their loss values as $\mathbb{V}_{\text{ave}}(m,n,k) = \{\boldsymbol{v}_{\text{ave},\,i}(m,n)|i = 1,\dots,k\}$ and $\mathbb{L}_{\text{ave}}(m,n,k) = \{L_{\text{ave},\,i}(m,n)|i = 1,\dots,k\}$, as well as the re-optimized solutions and their loss value as $\mathbb{V}_{\text{re-op}}(m,n,k) = \{\boldsymbol{v}_{\text{re-op},\,i}(m,n)|i = 1,\dots,k\}$ and $\mathbb{L}_{\text{re-op}}(m,n,k) = \{L_{\text{re-op},\,i}(m,n)|i = 1,\dots,k\}$. The distribution of all sets $\mathbb{L}_{\text{sel}}(20000,100,30)$, $\mathbb{L}_{\text{ave}}(20000,100,30)$ and $\mathbb{L}_{\text{re-op}}(20000,100,30)$ are compared in Figure 2D and 2H. The Euclidean distances between each solution type ($\boldsymbol{v}_{\text{sel}}$, $\boldsymbol{v}_{\text{ave}}$ and $\boldsymbol{v}_{\text{re-op}}$) and the raw predefined flux $\boldsymbol{v}'$ are calculated as $\mathbb{D}_{\text{sel, raw}}(m,n,k)$, $\mathbb{D}_{\text{ave, raw}}(m,n,k)$ and $\mathbb{D}_{\text{re-op, raw}}(m,n,k)$ by this equation:

$$\mathbb{D}_{(\cdot),\,\text{raw}}(m,n,k) = \left\{\left\|\boldsymbol{v}_{(\cdot)} - \boldsymbol{v}'\right\|_2 \middle| \forall \boldsymbol{v}_{(\cdot)} \in \mathbb{V}_{(\cdot)}(m,n,k)\right\} \tag{S19}$$

in which the $\|\cdot\|_2$ represents L2-norm of the target vector. Distribution of $\mathbb{D}_{\text{sel, raw}}(20000,100,30)$, $\mathbb{D}_{\text{ave, raw}}(20000,100,30)$ and $\mathbb{D}_{\text{re-op, raw}}(20000,100,30)$ are compared in Figure S2A and S2E.

Subsequently, for each selected solution in $\mathbb{V}_{\text{sel}}(m,n,k)$, average solution in $\mathbb{V}_{\text{ave}}(m,n,k)$ and re-optimized solution in $\mathbb{V}_{\text{re-op}}(m,n,k)$, their net flux vectors $\boldsymbol{v}_{\text{sel, net}}$, $\boldsymbol{v}_{\text{ave, net}}$ and $\boldsymbol{v}_{\text{re-op, net}}$ are computed based on Eq. S17 to $\mathbb{V}_{\text{sel, net}}(m,n,k)$, $\mathbb{V}_{\text{ave, net}}(m,n,k)$ and $\mathbb{V}_{\text{re-op, net}}(m,n,k)$. The direction of each reversible flux pair aligns with the same direction as in $\boldsymbol{v}'_{\text{net}}$. We compute the differential vectors between $\boldsymbol{v}'_{\text{net}}$ and each solution type, resulting in three vector sets $\mathbb{P}_{\text{sel}}(m,n,k)$, $\mathbb{P}_{\text{ave}}(m,n,k)$ and $\mathbb{P}_{\text{re-op}}(m,n,k)$. Euclidean distances and relative errors of $j$-th flux for each solution type to $\boldsymbol{v}'_{\text{net}}$ are also defined to form sets $\mathbb{D}_{\text{sel}}(m,n,k)$, $\mathbb{D}_{\text{ave}}(m,n,k)$ and $\mathbb{D}_{\text{re-op}}(m,n,k)$, $\mathbb{R}_{\text{sel},\,j}(m,n,k)$, $\mathbb{R}_{\text{ave},\,j}(m,n,k)$ and $\mathbb{R}_{\text{re-op},\,j}(m,n,k)$, respectively. The detailed equations are:

$$\mathbb{P}_{(\cdot)}(m,n,k) = \left\{\boldsymbol{v}_{(\cdot),\,\text{net}} - \boldsymbol{v}'_{\text{net}} \middle| \forall \boldsymbol{v}_{(\cdot),\,\text{net}} \in \mathbb{V}_{(\cdot),\,\text{net}}(m,n,k)\right\} \tag{S20}$$

$$\mathbb{D}_{(\cdot)}(m,n,k) = \left\{\left\|\boldsymbol{v}_{(\cdot),\,\text{net}} - \boldsymbol{v}'_{\text{net}}\right\|_2 \middle| \forall \boldsymbol{v}_{(\cdot),\,\text{net}} \in \mathbb{V}_{(\cdot),\,\text{net}}(m,n,k)\right\} \tag{S21}$$

$$\mathbb{R}_{(\cdot),j}(m,n,k) = \left\{\frac{v_{(\cdot),\,\text{net},j} - v'_{\text{net},j}}{v'_{\text{net},j} + \varepsilon} \middle| \forall \boldsymbol{v}_{(\cdot),\,\text{net}} \in \mathbb{V}_{(\cdot),\,\text{net}}(m,n,k)\right\} \tag{S22}$$

These net Euclidean distances are calculated as Eq. S18, in which the subtraction $\boldsymbol{v}_{(\cdot),\,\text{net}}(m,n) - \boldsymbol{v}'_{\text{net}}$ represents the differential vector of net fluxes and the $\varepsilon$ is a small number introduced for numeric stability. Distribution of $\mathbb{D}_{\text{sel}}(20000,100,30)$, $\mathbb{D}_{\text{ave}}(20000,100,30)$ and $\mathbb{D}_{\text{re-op}}(20000,100,30)$ are compared in Figure 2D and 2H. Distribution of $\mathbb{P}_{\text{sel}}(20000,100,30)$, $\mathbb{P}_{\text{ave}}(20000,100,30)$ and $\mathbb{P}_{\text{re-op}}(20000,100,30)$ are compared in Figure 2E and 2I. Distribution of $\mathbb{R}_{\text{sel}}(20000,100,30)$, $\mathbb{R}_{\text{ave}}(20000,100,30)$ and $\mathbb{R}_{\text{re-op}}(20000,100,30)$ are compared in Figure S2B and S2F.

As mentioned before, the degree of freedom of the metabolic network in our study is around 50, which means size of solution space is around $1000^{50} = 10^{150}$. Given the enormous size of the solution space, any reasonable value of $n$ would still be significantly smaller than the total solution space size. Therefore, the magnitude of $n$ is less likely to significantly affect the precision of the results but rather influences the stability of the averaged solutions. Conversely, selecting a smaller proportion of solutions from the total optimized solutions should theoretically yield results more aligned with the data, potentially enhancing the precision of averaged solutions.

To quantify the exact influence of parameters $m$ and $n$, we calculated these performance metrics under each combination of $m$ and $n$. For each combination of $m$ and $n$, comparison between $\mathbb{L}_{\text{sel}}(m, n, 30)$ and $\mathbb{L}_{\text{ave}}(m, n, 30)$, as well as between $\mathbb{D}_{\text{sel}}(m, n, 30)$ and $\mathbb{D}_{\text{ave}}(m, n, 30)$, sheds light on how the operations of selection and averaging impacts the loss value and the corresponding distance to the predefined flux vector as the parameters $m$ and $n$ varies. These analyses are presented in Figure S2D and S2H. For a more transparent visualization of each parameter's role, we introduce the selection ratio $m/n$. We calculated the average and the standard deviation (STD) of the set $\mathbb{D}_{\text{ave}}(m, n, 30)$ and depicted the dependencies of the average-to-selection ratio and the STD-to-$n$ in Figure 2F.

#### Robustness analysis to data limitations

In robustness analysis to data limitations, we first generate raw simulated MID data based on the predefined flux $\boldsymbol{v}'$. In the subsequent step, we shrink the data availability to generate a updated MID dataset for optimization. We then compare the optimized solution with the predefined flux to evaluate how these perturbations influence the precision and accuracy of MFA. To maintain clarity, we refer to the step involving the generation of simulated data as the “simulated step”, while the subsequent optimization with perturbed data availability is termed the “formal optimization”. In the formal optimization process, the optimization protocol closely follows that of the MFA protocol analysis, generating selected and averaged solutions $\mathbb{V}_{\text{sel, net}}(20000, 100, 5)$ and $\mathbb{V}_{\text{ave, net}}(20000, 100, 5)$. Their loss values $\mathbb{L}_{\text{sel}}(20000, 100, 5)$**,** $\mathbb{L}_{\text{ave}}(20000, 100, 5)$, as well as difference and relative error from the solution $\boldsymbol{v}'_{\text{net}}$ including $\mathbb{D}_{\text{sel}}(20000, 100, 5)$, $\mathbb{D}_{\text{ave}}(20000, 100, 5)$, $\mathbb{R}_{\text{sel}}(20000, 100, 5)$ and $\mathbb{R}_{\text{ave}}(20000, 100, 5)$, are calculated as per Eq. S21 and Eq. S22 to reflect the influence of these perturbations on data availability. These results are displayed in Figure 3 and S3.

In the perturbations on data availability, a constrained subset of metabolite MID in the generated simulated dataset is employed for subsequent optimization. This scenario aims to replicate the real-world situation where only a portion of metabolites can be measured by MS. There are three major perturbation styles in data availability: evenly distribution of MIDs, pathway-specific MID exclusion, and compartmental MID measurement. Within the “even distribution of limited MIDs” category, the perturbations labeled as “experimentally-available”, “medium”, and “few” represent different scenarios involving a varying number of available MIDs evenly distributed in all pathways. The “pathway-specific MIDs removal” perturbations signify situations where the MIDs from a specific pathway, denoted as “PPP”, “AA”, and “TCA”, are completely removed from the data set. The last “compartmental MID measurement” perturbations quantify the influence of whether the MID of the same metabolite located in different compartments can be measured separately.

#### Experimental data analysis

In this study, to minimize random errors stemming from batch preparation and measurement, the MIDs obtained from all biological replicates are averaged to create the target experimental MID for MFA. The MFA protocol used for the experimental data mirrors the one employed in the MFA protocol analysis with a fixed setting of $= 20000$ and $m = 100$. The net flux vector is computed for all 100 selected solutions using Eq. S17. This process is repeated for five times to obtain the set $\mathbb{V}_{\text{ave, net}}(20000{,}100{,}5)$, of which the mean is calculated as the final solution ($\mathbb{V}_{\text{ave, net}}(20000{,}100{,}3)$ in lung tumor dataset). Additionally, the standard deviation (STD) is determined to represent the uncertainty associated with the estimated average flux.

Since some datasets originate from infusion experiments involving patients, there are often limitations on infusion rates and durations due to safety considerations. Consequently, certain high-abundance metabolites, such as glucose (GLC_c) and citrate (CIT_m), may not become fully saturated by $^{13}C$ isotopomers, making it challenging to accurately fit their MIDs. To address this issue, an additional supplementary flux is introduced, assuming the presence of a pair of exchange fluxes between the unsaturated metabolite and an unlabeled pool:

$$\text{GLC\_stock} \underset{\text{GLC\_sup\_\_R}}{\overset{\text{GLC\_sup}}{\rightleftharpoons}} \text{GLC\_c} \tag{S23}$$

Here, "GLC_stock" represents the unlabeled pool with a fixed unlabeled MID. It is constrained that "GLC_sup" and "GLC_sup__R" are equal, meaning that this supplementary flux does not alter the current flux balance relationship. Instead, it provides extra flexibility to simulate the unsaturated labeling state.

In visualizing the flux solution, the corresponding main value and STD of the set $\mathbb{V}_{\text{ave, net}}(20000,100,5)$ are presented. To elucidate the relationships between different fluxes, various indices involving multiple fluxes are computed based on each flux vector in the set $\mathbb{V}_{\text{ave, net}}(20000,100,5)$. The corresponding mean and STD of all flux vectors are calculated for each index and visualized alongside other net fluxes. These indexes encompass:

$$\text{Lactate flux / Glycolysis flux: } \frac{\text{LDH\_c net}}{\text{GAPD\_c net}}$$

$$\text{TCA flux / Glycolysis flux: } \frac{\text{CS\_m}}{\text{GAPD\_c net}}$$

$$\text{Non-canonical / total TCA flux: } \frac{\text{CIT\_trans net}}{\text{CS\_m}}$$

$$\text{AMA / CMA: } \frac{\text{AKGMAL\_m net}}{\text{CIT\_trans net}}$$

#### Visualization of flux in network diagram

In all the figures, network diagrams, whether displaying flux values or not, are generated using the Python package Matplotlib. In the network diagrams with flux values, the directions of the flux arrows are determined by the directions of the corresponding net flux values, and the transparency of the flux arrows is based on the corresponding flux value. Since only a portion of the flux is depicted in the network diagrams, it is essential to clarify which flux value corresponds to each flux arrow:

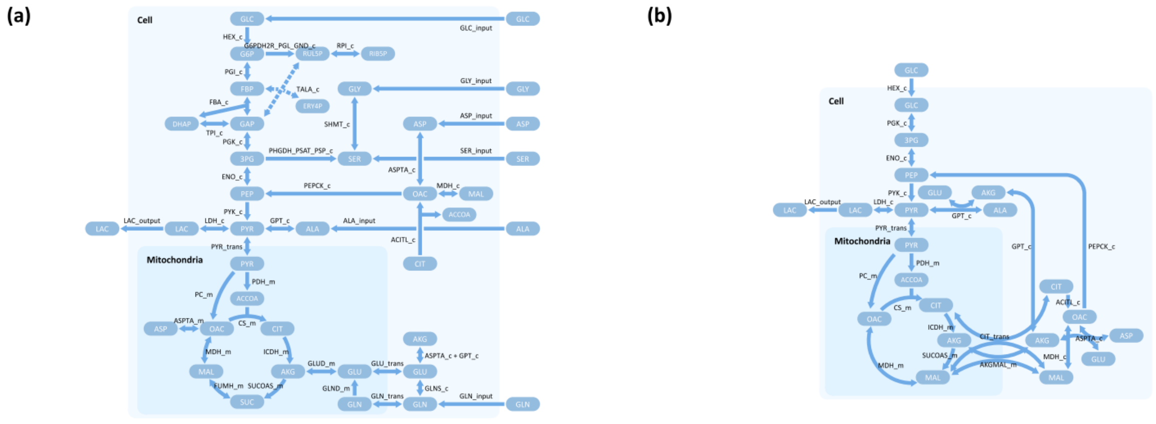

**Schematic 1**. Name of flux value that determined direction and transparency of flux arrow in normal (**a**) and exchange (**b**) metabolic network diagram

The transparency of flux arrows is calculated from the net flux value using a segmented linear function. This segmented linear function, denoted as $F_{SL}(x)$, is defined by a series of pairs $\{(x_i, y_i)|i = 1, \ldots, n\}$ and boundary $y$ value pair $(y_{\min}, y_{\max})$, ensuring that $F_{SL}(x)$ is a linear function within each segment, as follows:

$$F_{SL}(x) = \begin{cases} y_{\min} & x \leq x_1 \\ \frac{y_{i+1} - y_i}{x_{i+1} - x_i}(x - x_i) + y_i & x_i < x \leq x_{i+1} \\ y_{\max} & x > x_n \end{cases} \quad \text{(S24)}$$

In this study, the definition of $\{(x_i, y_i)|i = 1, \ldots, n\}$ is carefully designed to ensure that both $x_i$ and $y_i$ form monotonically increasing sequences. Consequently, $F_{SL}(x)$ is also monotonically increasing. Since the ultimate output represents transparency, the values of $y_{\min}$ and $y_{\max}$ are set to 0 and 1, respectively.

In the previously published HCT-116 data (Fig. 1), the defined sequence is $\{(0,0.05), (50, 0.55), (200, 0.8), (350, 1)\}$. This setting ensures a significant change in transparency within the flux value range $(0, 50)$, where most of the small fluxes are situated.

In the previously published human renal carcinoma data (Fig. 4), the defined sequence is $\{(0,0.03), (20, 0.3), (100, 0.75), (200, 0.9), (500, 1)\}$. This setting guarantees a notable change in transparency within the flux value range $(20, 100)$, where most of the fluxes are found.

In our new colon cancer data (Fig. 5), the defined sequence is $\{(0,0.03), (20, 0.4), (50, 0.65), (100, 0.7), (200, 0.9), (500, 1)\}$. This setting ensures a significant change in transparency in two specific flux value ranges, namely $(20, 50)$ and $(100, 200)$, where most of the fluxes are concentrated.

#### Software implementation

The scripts used in this study are developed in Python 3.8, and the corresponding source codes are accessible on GitHub at the following link: https://github.com/LocasaleLab/Scalable-automated-MFA-2023. Detailed information regarding package version dependencies can also be found on the GitHub website.

The computations are performed on a desktop PC equipped with either an i9-12900KS or i7-13700K CPU. To expedite the computation process, several strategies, such as parallel processing, have been employed. It's worth noting that each experiment typically consists of 10-20 datasets, necessitating approximately 200,000 to 400,000 optimizations under the parameter $n = 20000$. With each optimization taking roughly 2 seconds (Fig. 1D), completing one experiment generally requires between 100 to 200 hours of CPU time. Additionally, this duration increases proportionally with the number of repetitions needed in calculations (e.g., $k = 30$ in our algorithm development and $k = 5$ in our experimental dataset analyses).

##### Glucose starvation experiments in colon cancer cell lines

Our research incorporates glucose starvation experiments, a critical technique in cancer biology for studying tumor cell responses to glucose deprivation. This approach, involving exposing cancer cells to low-glucose conditions (below 1 mM), has been widely used in studies with various types of cancer cells, including colon cancer cells (such as HCT116[9], also used in our study), pancreatic cancer cells[10], and leukemia cells (such as Jurkat[11]). These studies consistently reveal that cancer cells can adapt to such limited glucose conditions, albeit with reduced growth rates and altered metabolic behaviors.

In line with these findings, we observed slower growth rates in our cancer cell lines, indicating cellular adaptation to restricted glucose availability. This suggests that limited nutrition prompts a significant yet predictable metabolic reprogramming, supporting the validity and relevance of conducting flux analysis based on these experimental conditions. Our results affirm that many metabolic fluxes decrease significantly under nutritional constraints (Figure 5D). Moreover, we quantified and analyzed ratios like CMA flux to TCA cycle flux and CMA flux to AMA flux (Figure 5E). These analyses highlight that, despite overall reductions in core metabolic fluxes, CMA flux decreases more markedly than other fluxes, including those related to AMA.

#### Model, data and configurations in each specific MFA

This section provides comprehensive information about each MFA experiment conducted in this study. For every MFA and simulated dataset, a set of three files is typically available, each containing specific details:

`solver_descriptions.xlsx`: This file contains detailed information about the MFA experiment, encompassing the model, data, and configurations used for MFA analysis.
`flux_raw_data.xlsx`: This file includes raw flux results obtained from the MFA computations.
`mid_raw_data.xlsx`: This file contains the predicted MID data of those raw flux results.

#### MFA for HCT116 cultured cancer cell line (figure 1, figure S1)

In this MFA analysis, the MID data used is from a previously published dataset[12]. To explore the solution space, 10000 random initial points are generated, and optimizations are performed independently from 400 of these random initial points. The best solution with the minimal loss value among 400 optimized results is plotted in the flux map. Furthermore, the net Euclidean distances between this best solution and other optimized solutions (including the No.2-5 best solutions, the top 20, and the top 100 optimized solutions) as well as random fluxes are calculated. To provide a comprehensive overview of distributions of the random fluxes and optimized solutions, the results from the 400 optimized solutions are embedded in a 2D space alongside the 400 random initial points.

The raw data folder for this MFA experiment is located at `raw_data/experimental_data_analysis/hct116_cultured_cell_line`. Detailed model, data and configurations of MFA, raw flux data and predicted MID data can be found in `solver_descriptions.xlsx`, `flux_raw_data.xlsx`, and `mid_raw_data.xlsx` in that folder respectively.

#### Simulated data

The predefined fluxes, as well as corresponding precise and noisy MID (both all-available and experimentally-available), in simulated data can be found in `raw_data/simulated_data/simulated_flux_vector_and_mid_data.xlsx` and `simulated_flux_vector_and_mid_data_with_noise.xlsx`.

#### Development and benchmark of the algorithm (figure 2, figure S2)

The MFA analysis in this study utilizes MID data generated from simulated data, which includes both all-available and experimentally-available datasets. To assess the sensitivity of the MFA results to different parameters, we vary the number of total optimizations ($n$) over the following values: $[100, 200, 500, 1000, 2000, 5000, 10000, 20000, 50000]$. Additionally, the number of selections ($m$) is tested using the following options: $[1, 2, 5, 10, 20, 50, 100, 200, 500]$.

The raw data folder for this MFA experiment is located at `raw_data/model_data_sensitivity`. Detailed raw flux data and predicted MID data of all-available and experimentally-available datasets can be found in the following folders: `raw_model_all_data` and `raw_model_raw_data` in that folder respectively.

#### Robustness analysis to data limitations (figure 3, figure S3)

In all scenarios of robustness analyses, the number of total optimizations ($n$) is set to 20,000, while the number of selections ($m$) is fixed at 100. Non-perturbed results in each sensitivity experiment are randomly selected from the raw flux data described in the "Protocol of MFA" section.

You can find the detailed model, data, configurations of MFA, raw flux data, and predicted MID data of data sensitivity experiments in the following folders: `raw_data/model_data_sensitivity/data_sensitivity`.

#### MFA for kidney, brain and lung tumor (figure 4, figure S4, figure S5)

The MID data used in this MFA analysis for patient infusion experiments is derived from previously published datasets[13]. As explained in the “Experimental data analysis” section, the models employed for these two datasets from infused patients incorporate two additional supplementary fluxes associated with GLC_c and CIT_m. In these analyses, the number of total optimizations ($n$) and the number of selections ($m$) remains fixed at 20,000 and 100 to calculate the average flux vector. The optimization-averaging process is repeated for five times (three times in lung tumor samples because of more samples in this experiment), of which the mean is calculated as the final solution and the standard deviation (STD) is determined to represent the uncertainty. These results are presented in Figure S5.

For detailed information on the models, data, configurations of MFA, raw flux data, and predicted MID data for kidney and brain tumor, please refer to the following folders: `raw_data/experimental_data_analysis/renal_carcinoma_invivo_infusion`

For lung tumor, please refer to the following folders:

`raw_data/experimental_data_analysis/lung_tumor_invivo_infusion`.

#### MFA for 8 cultured colon cancer cell lines (figure 5, figure S6)

The MID data used in this MFA analysis for the cultured cell lines were generated by our team. Similar to other experiments, the number of total optimizations ($n$) and the number of selections ($m$) remains fixed at 20,000 and 100 to calculate the average flux vector. The selection-averaging process is repeated for five times, of which the mean is calculated as the final solution and the standard deviation (STD) is determined to represent the uncertainty.

For detailed information on the models, data, configurations of MFA, raw flux data, and predicted MID data for the 8 cultured cancer cell lines, please consult the following folder:

`raw_data/experimental_data_analysis/colon_cancer_cell_line`.
